# Revealing virulence potential of clinical and environmental *Aspergillus fumigatus* isolates using Whole-genome sequencing

**DOI:** 10.1101/587402

**Authors:** F. Puértolas-Balint, J.W.A. Rossen, C. Oliveira dos Santos, M.A. Chlebowicz, E. Raangs, M.L. van Putten, P.J. Sola-Campoy, L. Han, M. Schmidt, S. García-Cobos

**Author notes:** Corresponding Author: S. García-Cobos;. Address: University of Groningen, University Medical Center Groningen, Department of Medical Microbiology and Infection Prevention EB80, Hanzeplein 1, P.O. Box 30.001, 9700 RB Groningen, The Netherlands.

## Abstract

*Aspergillus fumigatus* is an opportunistic airborne pathogen and one of the most common causative agents of human fungal infections. A restricted number of virulence factors have been described but none of them lead to a differentiation of the virulence level among different strains. In this study, we analyzed the whole-genome sequence of a set of *A. fumigatus* isolates from clinical and environmental origin to compare their genomes and to determine their virulence profiles. For this purpose, a database containing 244 genes known to be associated with virulence was built. The genes were classified according to their biological function into factors involved in thermotolerance, resistance to immune responses, cell wall structure, toxins and secondary metabolites, allergens, nutrient uptake and signaling and regulation. No difference in virulence profiles was found between clinical isolates causing an infection and a colonizing clinical isolate, nor between isolates from clinical and environmental origin. We observed the presence of genetic repetitive elements located next to virulence related gene groups, which could potentially influence their regulation. In conclusion, our genomic analysis reveals that *A. fumigatus*, independently of their source of isolation, are potentially pathogenic at the genomic level, which may lead to fatal infections in vulnerable patients. However, other determinants such as genetic variations in virulence related genes and host-pathogen interactions most likely influence *A. fumigatus* pathogenicity and further studies should be performed.

**Importance:** *Aspergillus* spp. infections are among the most clinically relevant fungal infections also presenting treatment difficulties due to increasing antifungal resistance. The lack of key virulence factors and a broad genomic diversity complicates the development of targeted diagnosis and novel treatment strategies. A widely spread variability in virulence has been reported for experimental, clinical and environmental isolates. Here we provide supporting evidence that members of this species are fully capable of establishing an infection in immunosuppressed hosts according to their virulence content at the genomic level. Due to the possible clinical complications, studies are urgently required linking strain’s virulent phenotype with the genotype to better understand the virulence activation of this important fungal pathogen.

## Introduction

*Aspergillus fumigatus* (*A. fumigatus*) is an opportunistic fungal pathogen that poses one of the major threats to immunocompromised individuals in the clinic. High risk patients include neutropenic patients, hematopoietic stem cell transplant recipients, patients receiving a prolonged steroid treatment and critically-ill patient in the intensive care unit (ICU) with chronic obstructive pulmonary disease (COPD), liver cirrhosis, viral infections or microbial sepsis (1–3). In the context of an impaired immune function, inhaled airborne spores of *A. fumigatus* will not be effectively eliminated and will remain in human airways to cause a range of infections that include allergic bronchopulmonary aspergillosis (ABPA), aspergilloma (chronic aspergillosis) and invasive aspergillosis (IA) (1,4). IA is the most serious infection, with a global prevalence of 250,000 cases per year with mortality rates up to 90-95% (5,6). In addition to the increasing burden of patients with impaired immunity (1), another major challenge is the treatment of these fungal infections due to the antifungal resistance to triazoles, the most indicated drugs against *Aspergillus* species infections. Resistance is characterized by the presence of a point mutation (L98H) in the azole target *Cyp51A* and a 34-base pair (bp) tandem repeat (TR34) in its promoter region (7) and the most common cause of resistance acquisition is the widespread azole-based fungicide use against fungal plant pathogens in the agricultural practice (7–9).

In order to overcome the therapeutic challenges and threats posed by *A. fumigatus* infections, there is a need to further understand the mechanisms of adaptability and infection of the fungus to develop better and early diagnostic tools and uncover novel therapeutic strategies. The virulence of *A. fumigatus* is multifactorial, a trait that has been developed by the fungus as a need to survive the encountered selective pressures in decaying vegetation (10). Whole-genome and transcriptome analysis have allowed the discovery and study of new components of *A. fumigatus* biology and pathogenesis, providing a better understanding of the genetic content of *Aspergillus spp*. Genomic analyses identified that *A. fumigatus* contains 8.5% of lineage-specific (LS) genes with accessory functions for carbohydrate and amino acid metabolism, transport, detoxification, or secondary metabolite biosynthesis, suggesting this microorganism has particular genetic determinants that can facilitate an *in vivo* infection (11). Nevertheless, the study of *A. fumigatus* virulence has been hampered by the lack of a standard wild-type (WT) strain, next to a broad isolate-dependent variability in virulence as shown in murine infection models of IA (12). *A. fumigatus* isolates can be divided in three different categories depending on the source of isolation: 1) environmental, obtained from decaying vegetation, air sampling, crops, etc.; 2) clinical, originally found in patient samples, and 3) experimental, which refers to isolates that were first obtained from a patient setting but are now used as reference strains by many research groups (i.e. Af293 or CEA10). Some experimental infection studies have reported clinical isolates to be more virulent than environmental isolates (13–15). Moreover, two environmental isolates retrieved from an International Space Station were more virulent than the experimental strains Af293 and CEA10 (16). Additionally, an in-host study that analyzed the microevolution of 13 isogenic isolates of *A. fumigatus* obtained from the same patient over a period of 2 years, reported both increases and decreases in virulence among the isolates (17). The latter, most likely as a result of the adaptation of this microorganism to the human niche to allow its persistence (17). These different observations highlight the need to recognize the intraspecies genotypic and phenotypic variety among *A. fumigatus* populations, as this could determine the progression and fulminant outcome of diseases produced by this fungus.

To increase the knowledge on molecular factors contributing to the development of diseases caused by *A. fumigatus*, the present study aimed to determine the underlying genetic traits that characterize a virulent strain. We evaluated if differences in the afore mentioned strain-specific virulence can be explained through abundance of virulence factors at the genomic level, with special interest in clinical isolates obtained from patients with different clinical outcomes. A database containing 244 *A. fumigatus* virulence related genes (VRGs), reported by different studies, was created and used to define the pathogenic potential of investigated isolates. In total nine *A. fumigatus* sequences were used in our study: two experimental, five clinical and two environmental isolates. Among these isolates, the whole-genome sequences of three clinical isolates and one experimental strain B5233 were generated at the University Medical Center Groningen (UMCG), and further used for strain genotyping and to perform a comparative genomic analysis to identify genomic differences.

## Results

### Virulence related genes screening showed that all *A. fumigatus* isolates included in this study are potentially pathogenic

The genome sequences of nine *A. fumigatus* isolates (Table 1) were screened for the presence of particular VRGs using our in-house database (see Table S2). We identified the presence of all 244 VRGs (>90% coverage and >90% identity) in the genome of seven isolates P1MR, P1MS, P2CS, Af293, 12-7505054, 08-19-02-30 and 08-19-02-46. In addition, 243 genes were present in the genomes of B5233 and 08-12-12-13, and both isolates lacked the *Afu5g12720* gene. This gene codes for a putative ABC transporter and is a member of the Biosynthetic gene cluster 17 (BGC17), consisting of in total 10 genes, described by Bignell *et al* (18). The product of this BGC is a non-ribosomal peptide synthetase (NRPS), which is thought to be structural (19). However, no clear link between this ABC transporter and the function of this NRPS has been described before, thus it is not known how its absence could affect the overall function of this cluster and its possible role in virulence.

**TABLE 1.**
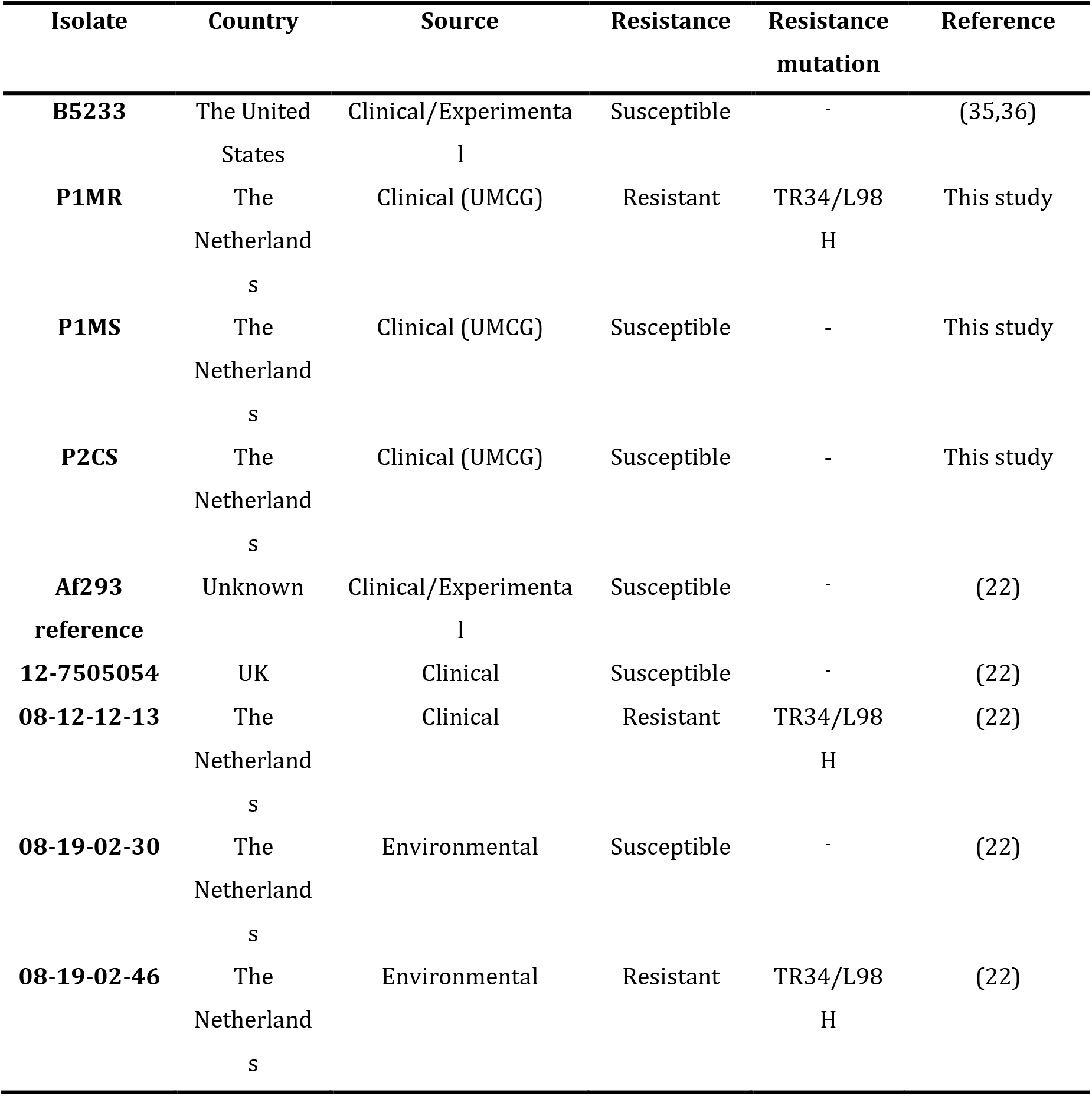
Characteristics of *A. fumigatus* isolates investigated in this study.

| Isolate | Country | Source | Resistance | Resistance mutation | Reference |
| --- | --- | --- | --- | --- | --- |
| **B5233** | The United States | Clinical/Experimental | Susceptible | - | (35,36) |
| **P1MR** | The Netherlands | Clinical (UMCG) | Resistant | TR34/L98 H | This study |
| **P1MS** | The Netherlands | Clinical (UMCG) | Susceptible | - | This study |
| **P2CS** | The Netherlands | Clinical (UMCG) | Susceptible | - | This study |
| **Af293 reference** | Unknown | Clinical/Experimental | Susceptible | - | (22) |
| **12-7505054** | UK | Clinical | Susceptible | - | (22) |
| **08-12-12-13** | The Netherlands | Clinical | Resistant | TR34/L98 H | (22) |
| **08-19-02-30** | The Netherlands | Environmental | Susceptible | - | (22) |
| **08-19-02-46** | The Netherlands | Environmental | Resistant | TR34/L98 H | (22) |

### TRESP genotyping identified a distinct genetic background of the isolates

Out of four investigated isolates, two were isolated from the same patient suffering from an influenza A H1N1 infection with IA (Figure 1). Given that P1MR, a resistant strain, was isolated nine days after a susceptible isolate P1MS was obtained, the question arose as to whether these isolates were genetically related and if the resistant phenotype developed after azole treatment. To determine their genetic relatedness, we used the recently described TRESP method that identifies the alleles of specific tandem repeats in the exon sequence of surface proteins CSP, MP2 and CFEM (Table 2) (20). The four isolates presented different allelic combinations and thus, different TRESP genotypes, P1MS and P1MR having t03m1.1c08A and t11m1.1c09 TRESP genotypes, respectively. In this study, CSP alleles appeared to best differentiate the isolates (Table 2).

**FIG 1.**
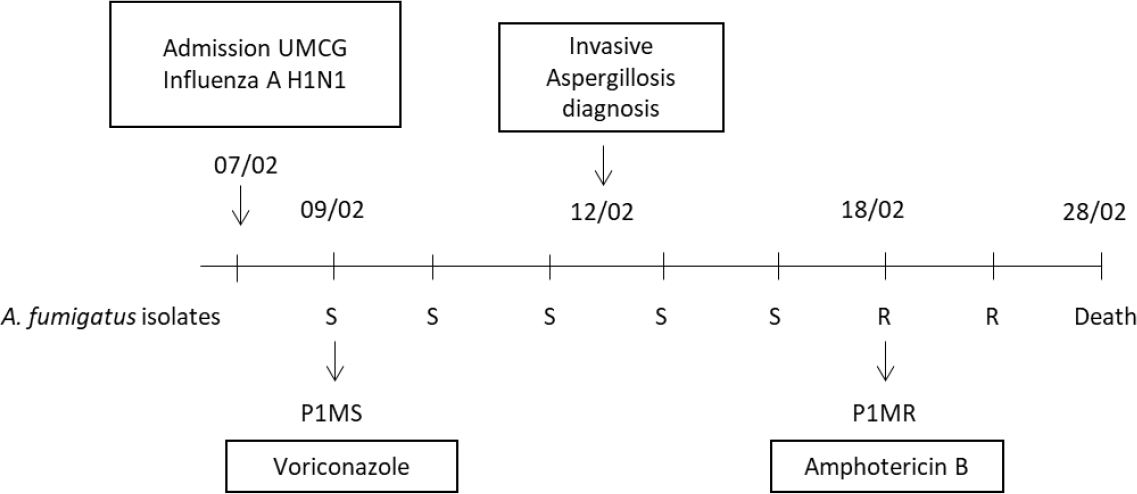
Time line of the influenza A H1N1 patient staying in the hospital and the course of infection. A total of seven *A. fumigatus* isolates were obtained from sputum samples, five susceptible (S) and two resistant (R) ones. The patient remained in the hospital for a period of 21 days until the time of death. Isolates P1MS and P1MR included in this study are indicated in the figure.

**TABLE 2.** TRESP genotype based on repetitive sequences in the exons of surface proteins CSP, MP2 and CFEM.

| Sample | Allele CSP | Allele MP2 | Allele CFEM | TRESP genotype |
| --- | --- | --- | --- | --- |
| B5233 | t02 | m1.2 | c09 | t02m1.1c09 |
| P1MR | t11 | m1.1 | c09 | t11m1.1c09 |
| P1MS | t03 | m1.1 | c08A | t03m1.1c08A |
| P2CS | t02 | m1.1 | c19 | t02m1.1c19 |

### Comparative genomics illustrates differences in the genome structure

The genomes of our four isolates were compared to *A. fumigatus* Af293 chromosomes. The compared genomic sequences of the 8 chromosomes are displayed in Figure 2, in which investigated VRGs locations are highlighted in yellow. Small deletions (100 kpb) were observed at the end of chromosomes 5 and 6, and large deletions (>300 kbp) were identified at the beginning of chromosome 1 and at the end of chromosome 7. Multiple small deletions and large-scale deletions in *A. fumigatus* genomes have been reported previously, and particularly these >300kbp large-scale deletions were previously described in chromosome 1 (21, 22) and chromosome 7 (21). A region with a high dissimilarly with respect to the reference Af293 between 1,698 kbp and 2,058 kbp is observed in chromosome 7 for all the isolates, where only the P2CS isolate had a certain degree of similarity (Figure 2). In addition, sequence gaps with no assigned CDS, (indicated by a red line in Figure 2), represent putative centromeres in all chromosomes and in the case of chromosome 4, a region of ribosomal DNA (represented by a dark blue line) (11). Notably, repeat rich sequence areas were identified in chromosomes 1, 2, 4, 6 and 8. These areas can be seen in Figure 2 by the alignment of many small contigs which coincides with a low GC content. In the case of chromosome 4, a group of virulence genes appears to be flanked by these repetitive regions on both sides, whereas some groups are only flanked on one side as depicted in chromosomes 6 and 8.

**FIG 2.**
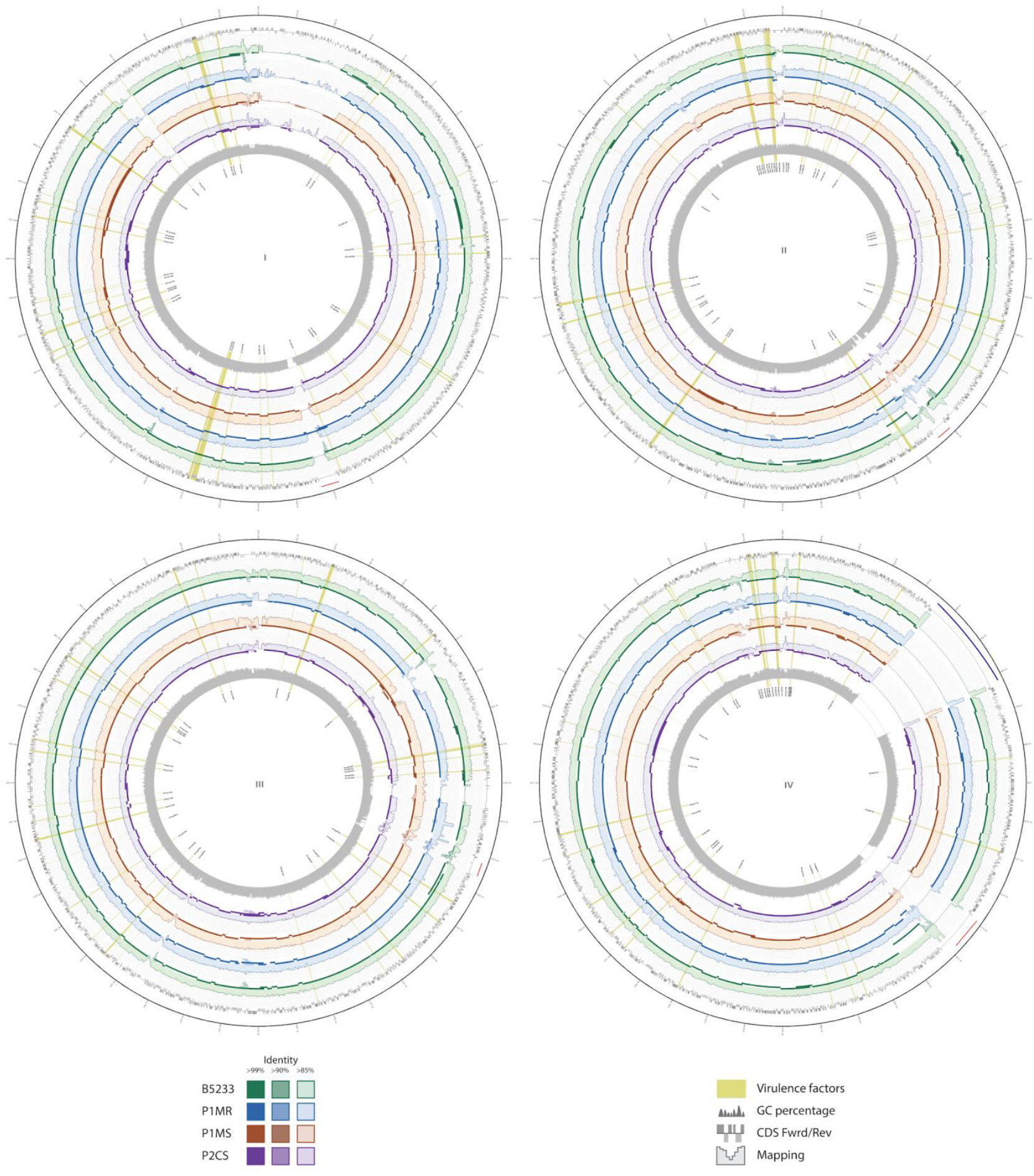
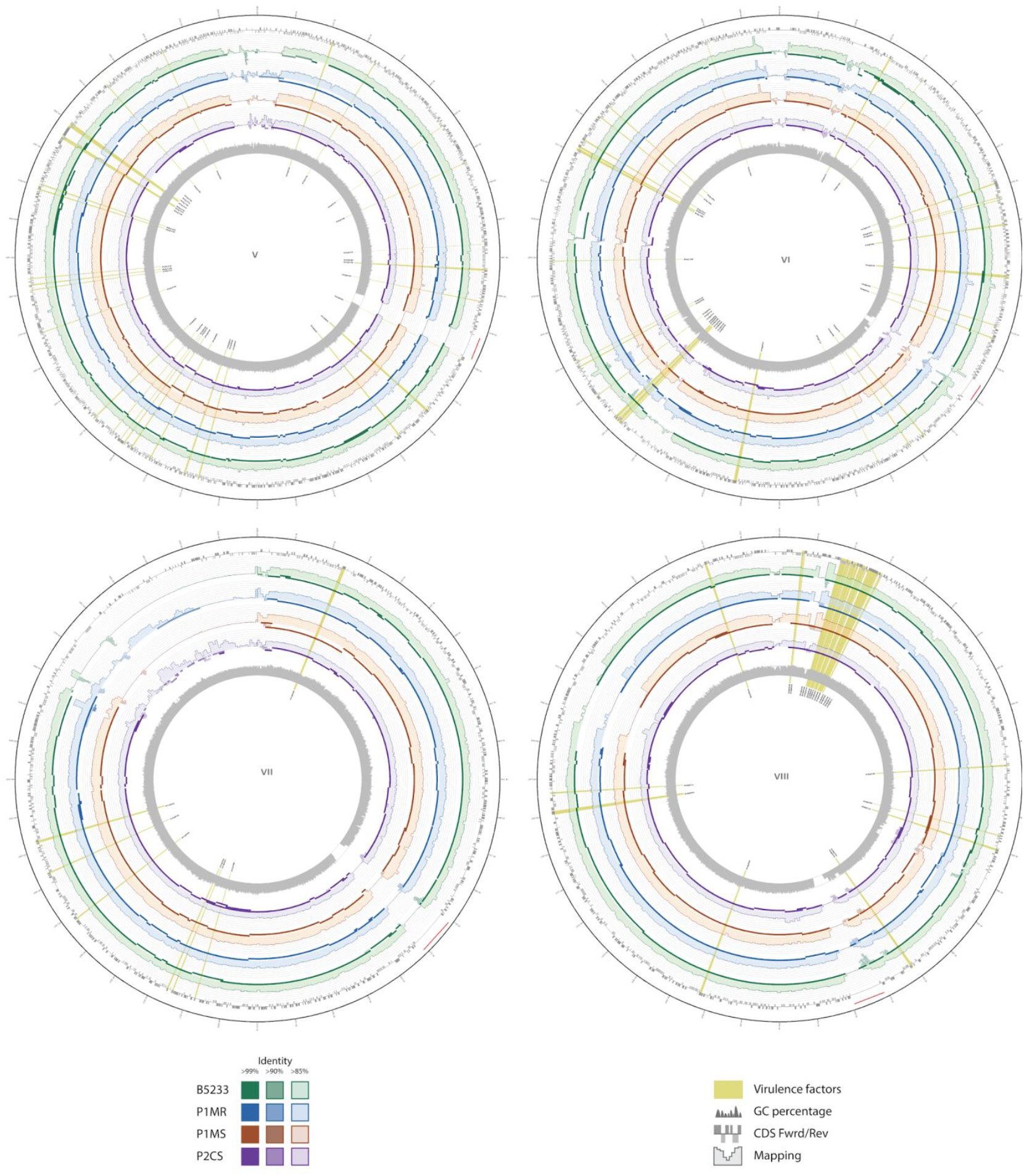
Graphical representation of assemblies and reads of B5233 (green), P1MR (blue), P1MS (orange) and P2CS (purple) isolates aligned to Af293 reference chromosomes (I, II, II, IV, V, VI, VII, VIII). Outer track indicates all CDS in forward (dark grey) or reverse (light grey) strand. Two different tracks are represented per isolate: one corresponding the mapping coverage and another one corresponding to contig alignment (minimum ID 85%). The complete contigs are represented with transparency in accordance to the local alignment identity. Genes related to virulence are highlighted in yellow with its names in the innermost track. GC% is represented every 100 bp. Red lines indicate putative centromeres and the dark blue line (chromosome 4) represents ribosomal DNA.

### Genomic variability among the fungal genomes

Variant calling using as reference the *A. fumigatus* Af293 genome, identified a total number of 68,352; 48,590; 56,362 and 56,422 variants in the genome of B5233, P1MS, P1MR and P2CS isolates, respectively (Table 3). High and moderate impact variants were retrieved, and their predicted effect is displayed in Table S3. Among variations with a predicted moderate- and high-impact, a high number of missense variants, ranging from 9,804 to 12,067, was identified. Single nucleotide polymorphism (SNP) analysis in VRGs with respect to reference Af293 revealed the presence of a range of 1,015 – 1,122 SNPs in all the analyzed isolates (Table 3). Examples of some variants present in the VRGs are listed in Table 4 and a more detailed description is given in supplementary Table S4. No clear pattern of variant distribution was found regarding the source the isolates. Rather, we observed some cases where all isolates had common variants as demonstrated in genes *thtA, sidC* and *msdS*. Genes associated to resistance to the immune response *rodB, cat1* and *afpmt2* had only one or no variants suggesting they are highly conserved genes. The *gliZ gene* required for the regulation of gliotoxin and the *gli* cluster, and the *sidC* gene, with an important role in iron acquisition, are examples of genes with several variants.

**TABLE 3.** Variant analysis of the novel *A. fumigatus* isolates against reference *A. fumigatus* Af293.

| Isolates | SnpEff | SnpSift Filter |  | SNPs present in the<br>virulence genes (CLC<br>Genomics Workbench) |
| --- | --- | --- | --- | --- |
|  | Total number<br>of variants | MODERATE | HIGH |  |
| B5233 | 68,352 | 12,085 | 884 | 1122 |
| P1MS | 48,590 | 10,109 | 752 | 1107 |
| P1MR | 56,362 | 11,718 | 870 | 1015 |
| P2CS | 56,422 | 12,085 | 884 | 1158 |

**TABLE 4.**
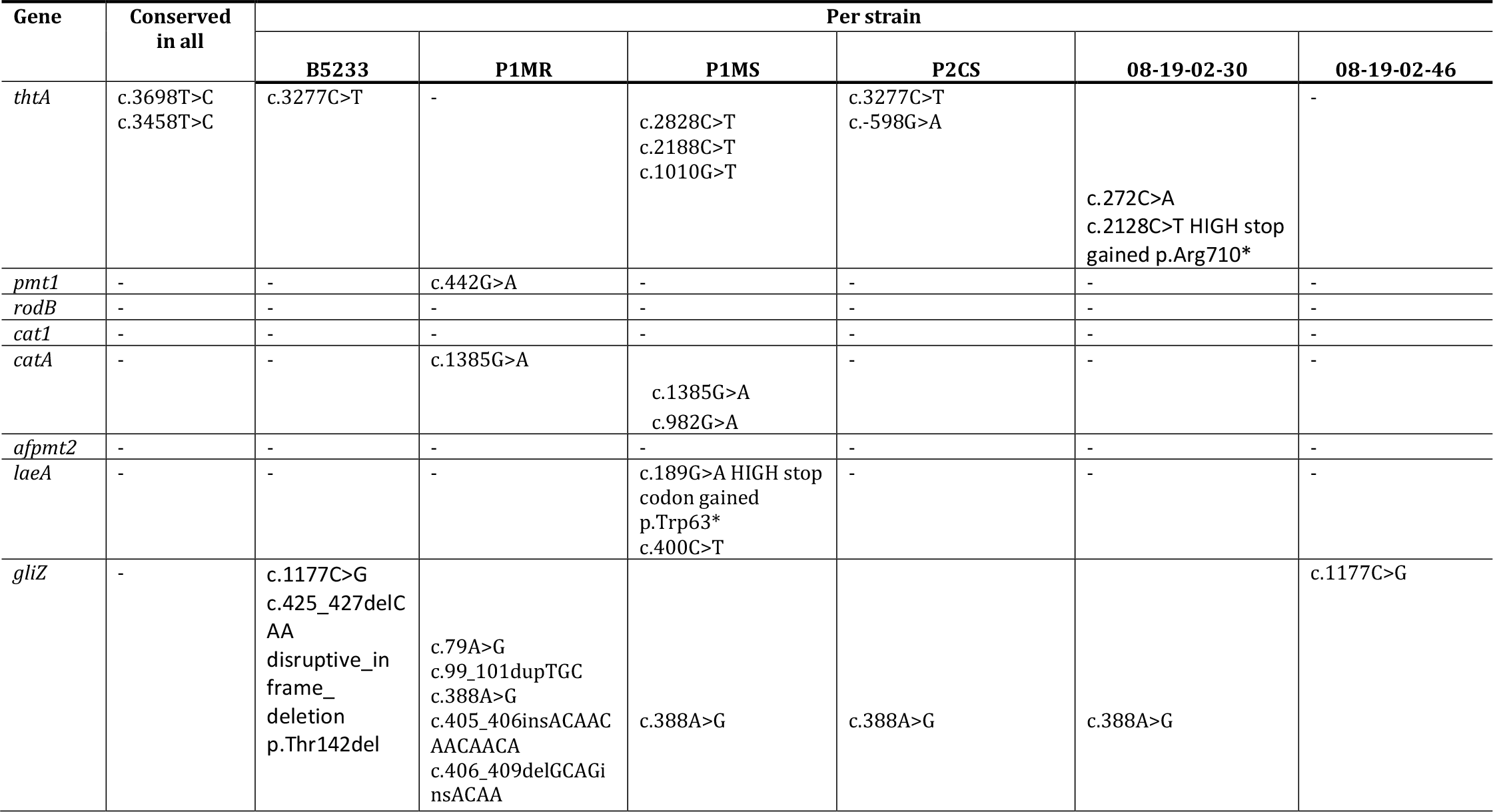
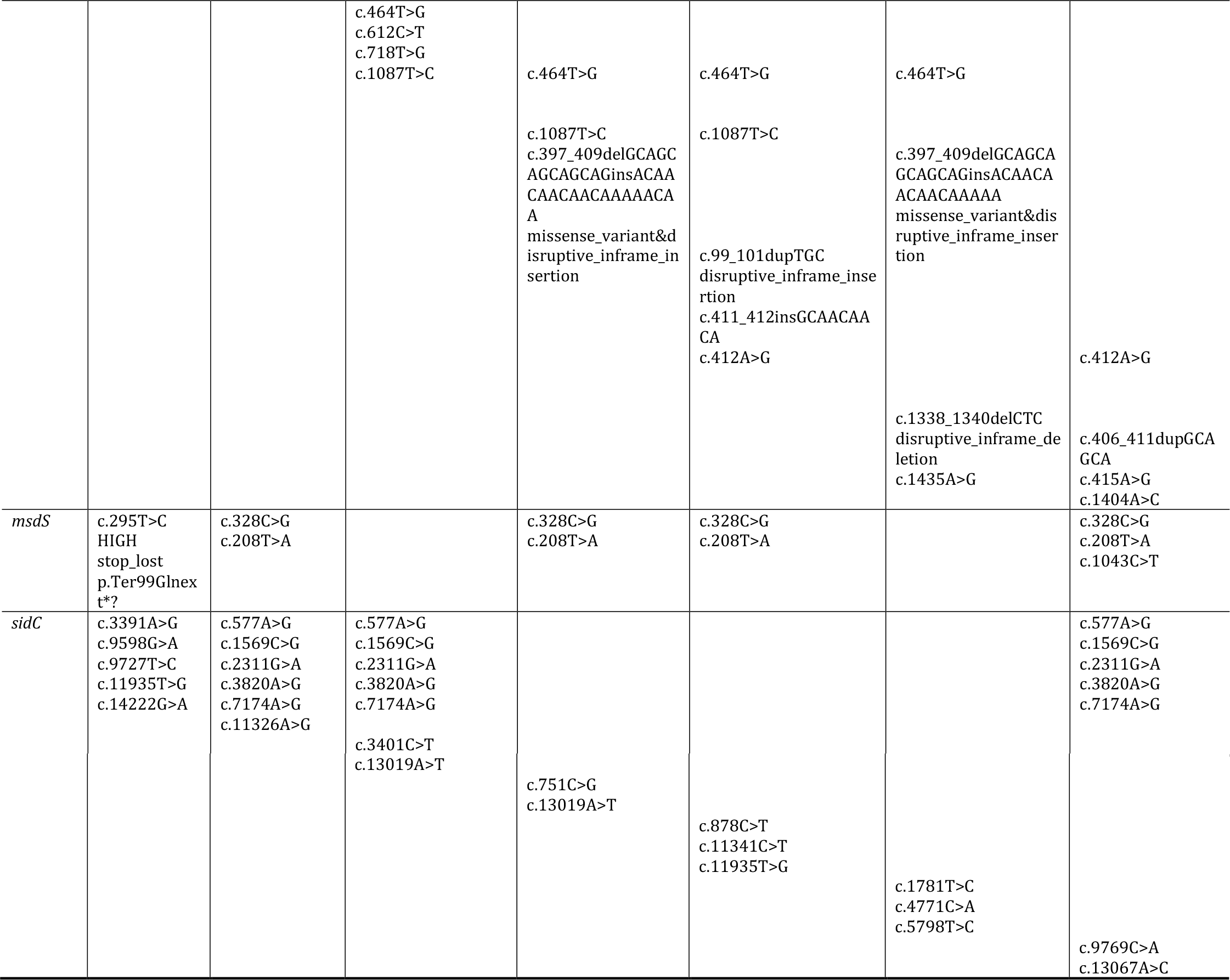
Examples of shared and unique moderate and high impact variants in genes associated to thermotolerance, resistance to the immune response, toxins and secondary metabolites, allergens and nutrient uptake.

| Gene | Conserved in all | Per strain |  |  |  |  |  |
| --- | --- | --- | --- | --- | --- | --- | --- |
|  |  | B5233 | P1MR | P1MS | P2CS | 08-19-02-30 | 08-19-02-46 |
| *thtA* | c.3698T>C<br>c.3458T>C | c.3277C>T | - | c.2828C>T<br>c.2188C>T<br>c.1010G>T | c.3277C>T<br>c.-598G>A | c.272C>A<br>c.2128C>T HIGH stop<br>gained p.Arg710* | - |
| *pmt1* | - | - | c.442G>A | - | - | - | - |
| *rodB* | - | - | - | - | - | - | - |
| *cat1* | - | - | - | - | - | - | - |
| *catA* | - | - | c.1385G>A | c.1385G>A<br>c.982G>A | - | - | - |
| *afpmt2* | - | - | - | - | - | - | - |
| *laeA* | - | - | - | c.189G>A HIGH stop<br>codon gained<br>p.Trp63*<br>c.400C>T | - | - | - |
| *gliZ* | - | c.1177C>G<br>c.425_427delC<br>AA<br>disruptive_in<br>frame_<br>deletion<br>p.Thr142del | c.79A>G<br>c.99_101dupTGC<br>c.388A>G<br>c.405_406insACAAC<br>AACAACA<br>c.406_409delGCAGi<br>nsACAA | c.388A>G | c.388A>G | c.388A>G | c.1177C>G |
|  |  |  | c.464T>G<br>c.612C>T<br>c.718T>G<br>c.1087T>C | c.464T>G<br>c.1087T>C<br>c.397_409delGCAGC<br>AGCAGCAGinsACAA<br>CAACAACAAAAACA<br>A<br>missense_variant&d<br>isruptive_inframe_in<br>sertion | c.464T>G<br>c.1087T>C<br>c.99_101dupTGC<br>disruptive_inframe_inse<br>rtion<br>c.411_412insGCAACAA<br>CA<br>c.412A>G | c.464T>G<br>c.397_409delGCAGCA<br>GCAGCAGinsACAACA<br>ACAACAAAAA<br>missense_variant&dis<br>ruptive_inframe_inser<br>tion<br>c.1338_1340delCTC<br>disruptive_inframe_de<br>letion<br>c.1435A>G | c.412A>G<br>c.406_411dupGCA<br>GCA<br>c.415A>G<br>c.1404A>C |
| *msdS* | c.295T>C<br>HIGH<br>stop_lost<br>p.Ter99Glnex<br>t*? | c.328C>G<br>c.208T>A |  | c.328C>G<br>c.208T>A | c.328C>G<br>c.208T>A |  | c.328C>G<br>c.208T>A<br>c.1043C>T |
| *sidC* | c.3391A>G<br>c.9598G>A<br>c.9727T>C<br>c.11935T>G<br>c.14222G>A | c.577A>G<br>c.1569C>G<br>c.2311G>A<br>c.3820A>G<br>c.7174A>G<br>c.11326A>G | c.577A>G<br>c.1569C>G<br>c.2311G>A<br>c.3820A>G<br>c.7174A>G<br>c.3401C>T |  |  |  | c.577A>G<br>c.1569C>G<br>c.2311G>A<br>c.3820A>G<br>c.7174A>G |
|  |  |  | c.13019A>T | c.751C>G<br>c.13019A>T | c.878C>T<br>c.11341C>T<br>c.11935T>G | c.1781T>C<br>c.4771C>A<br>c.5798T>C | c.9769C>A<br>c.13067A>C |

Additionally, a comparative analysis between P1MS and P1MR isolates from the same patient was performed and 45,335 variants were found, corresponding to 38,319 SNPs, 868 multiple nucleotide polymorphisms (MNPs), 1,768 insertions, 1,842 deletions and 2,538 complex (combination of a SNPs and MNPs).

## Discussion

*A. fumigatus* is a major fungal opportunistic pathogen capable of causing chronic and deadly invasive infections. The development of an infection seems to be primarily determined by the host immune status. Here, we performed a genomic analysis to investigate the role of the pathogen in the development and progression of the infection. We hypothesized that *A. fumigatus* isolates recovered from a patient who died after an infection with influenza A H1N1 and IA and, an isolate from a patient with human immunodeficiency virus (HIV) and COPD with no reported Aspergillus infection, would reveal a distinct virulence profile. In addition to our clinical isolates, we studied the known virulent *A. fumigatus* experimental strain B5233 and five unrelated isolates available in public databases with different sources of isolation (Table 1). Our analysis identified the presence of 244 VRGs in all *A. fumigatus* isolates tested, with the exception of *Afu5g12720* gene in B5233 and 08-12-12-13, indicating all isolates have the potential to be or become pathogenic. Thus, differences in the microorganisms’ capacity to cause damage most likely do not only rely on the presence or absence of virulence factors at the genomic level of *A. fumigatus*. Moreover, a high variability in the studied *A. fumigatus* genomes, reflects the enormous capacity of adaptation of the fungus to different environments.

The only gene amongst the 244 VRGs included in our in-house database, not found in the isolates B5233 and 08-12-12-13 was *Afu5g12720*. Interestingly, this gene was reported to be absent in 21 out of 66 *A. fumigatus* samples in a population genomics study that investigated the genomic variation of secondary metabolites in this species (23). This gene is a member of the BGC 17, and its absence could have a functional impact on the synthesis of the final product of this cluster, a NRPS, which is thought to have a structural function (19). *Afu5g12720* codes for an ABC transporter located in the BGC17 with another 9 genes (18) and was curiously absent in B5233, a strain that has been described as highly virulent. It would be interesting to further study the link between the lack of this gene and a possible increase in virulence, since disruption of another gene member included in this BGC17, *pes3*, resulted in a hypervirulent strain (19).

Additional changes in the genome structure of our isolates were found in our comparative genomic analysis. We identified deleted segments in relation to the reference that were located at the beginning of chromosome 1 and at the end of chromosomes 5, 6 and 7. Fedorova *et al*, identified these subtelomeric regions to be enriched for the presence of pseudogenes, transposons and other repetitive elements, and corresponding to areas that contain genes specific to the *A. fumigatus* species. (11) It was hypothesized that these genes have most likely evolved from big duplication and diversification events and not horizontal gene transfer. (11) It seems isolates B5233, P1MS and P1MR also do not have these genes. Our hypothesis is that these segments, are insertion-prone regions that are contributing to the diversification of the species.

Nucleotide variant analysis of our four isolates, identified a total number of variations ranging from 48,590 to 68,352 compared to reference strain Af293. This range fits in the genetic diversity for *A. fumigatus* reported previously and determined in 95 sequences ranging from 36 - 72,000 SNPs (16). The large number of identified variants and differences in the genome structure displays a broad genetic diversity in the studied isolates. This diversity is hypothesized to directly influence the fungus virulence by allowing an adaptation to an in-host environment, the evasion of the host immune system and the acquisition of antifungal resistance (17,24,25,27). The presence of SNPs in the VRGs of the clinical isolates, particularly those to be predicted to have a high impact, could be of major influence for the virulence of the isolates. However, in this study we could not link the presence of genetic changes in VRGs to isolates with a common origin of isolation. In addition, some repetitive elements were located on the sides of some groups of VRG as exemplified on chromosomes 6 and 8. It is possible they could play a role in how some of these genes are expressed, since these elements are recognized to shape the genomes of fungi (28). Follow-up studies using RNA sequencing could help to elucidate how and under what circumstances these virulence genes are expressed, as well as to determine the impact of genomic variations on expression levels. Subsequent infection model studies could be used to correlate these genomic variations and changes with specific pathogenic phenotypes.

The genome sequence of isolates P1MS and P1MR differed by 45,335 variants and they were confirmed to have different genotypes, supporting the hypothesis that resistance did not develop from the initial susceptible isolate. It is also unlikely that in a period of nine days the susceptible isolate would have been able to mutate to acquire azole resistance since the median time of development of azole resistance has been reported to be 4 months (29). Moreover, gain of the resistant phenotype within the host is observed in chronic infections whereas acquisition of resistance during IA continues to be unreported (24). In a similar case of post-influenza aspergillosis, four *A. fumigatus* isolates were obtained from a patient that received an allogeneic stem cell transplant and developed IA after the influenza virus infection which was initially treated with voriconazole (30). The patient passed away 4 months later. The first three isolates were susceptible to azole treatment but the last one, demonstrated triazole-resistance. It was confirmed that the resistant isolate was different from the first isolates by STRA*f* microsatellite genotyping (30). It is likely that the resistant phenotype of the resistant isolate in both our study and the post-influenza study (30), was of environmental origin and that this isolate had coexisted with the susceptible isolates in a mixed population that was not detected during the first sampling. Treatment with voriconazole most probably eradicated the initial susceptible strain and through selective pressure, allowed the resistant *A. fumigatus* strains to persist in the patient’s airways. Because of this possibility, a change of practice regarding *A. fumigatus* isolation has been applied at the diagnostics laboratory at the UMCG where antifungal susceptibility testing is now applied to at least five colonies obtained from a single respiratory sample. Influenza virus infection has been recently described as an independent risk factor for invasive pulmonary aspergillosis and, therefore, extreme care for patients admitted into the ICU with a severe influenza virus infection is advised (31).

The use of TRESP genotyping in this study identified the isolates to be genetically different. This approach was easy and accessible and only required the whole-genome sequence of the isolates, in contrast to other traditional typing methods with much lower discriminatory power (MLST) (32), laborious microsatellite determination (STRA*f*) (33), and the most novel whole-genome SNP based typing method which is highly dependent on the quality of sequencing, variant calling parameters and selection of a genetically close reference strain (21).

We show evidence that supports the observation that all fungi have the ability to cause disease and that members of the *A. fumigatus* species lack the sophisticated virulence factors, commonly used to describe differences in virulence in species of the bacterial kingdom, that could explain differences in their pathogenic traits (25,34). In order to define what makes a virulent *A. fumigatus* isolate many researchers have attempted characterizing different aspects of the fungus: differences in the colonial and spore color phenotype (25), the strain-dependent immunomodulatory properties induced in the host (25), the clinical or environmental source of the isolate (13, 25,26), the strains ability to adapt and grow in stressful conditions like low oxygen microenvironments where hypoxia fitness was strongly correlated with an increase in virulence (26), and the ability of the fungus to adjust its gene expression to survive in different immunosuppressive conditions inside the host (3). These are all aspects that influence how fit a strain will be to produce an infection and further research on the virulence of this microorganism should take all these aspects into consideration when characterizing an experimental and/or clinical strain. The combined results of these studies could be used to explore the link between the virulent phenotype and genotype to better understand the mechanisms of infection of this important human pathogen.

There are some limitations to be considered in this study. First, the number of isolates was small, however three different *A. fumigatus* population sources (clinical, environmental and experimental) were included. Nevertheless, we encourage our findings to be confirmed in a larger population to fully confirm the observation that all members of this species are potentially pathogenic. Second, we included 244 genes in our in-house database and we do not rule out the possibility that other genes that have not yet been characterized/described could also be considered as VRG.

In conclusion, we developed an in-house database with 244 VRG and found them all (except *Afu5g12720*) in the whole-genome sequence of nine *A. fumigatus* isolates, five clinical, two environmental and two experimental. This indicates, the strain-specific virulence genomic profile cannot be explained by the source of isolation when looking at differences in virulence related gene content or sequence variations. Understanding under what circumstances VRGs are expressed and utilized may ultimately contribute to explain how they regulate their virulence. Moreover, a broad genomic variability and the convenient location of transposable elements that are acknowledged to be shaping the genome, evidences a very efficient capacity of adaptation and challenges the development of specific diagnostic tools and effective treatments.

## Methods

### *A. fumigatus* isolates and background

*A. fumigatus* samples evaluated in this study are summarized in Table 1. Four clinical isolates were included: three isolates (P1MS, P1MR and P2CS) obtained at the University Medical Center Groningen (UMCG), Groningen, the Netherlands, and the strain B5233, kindly provided by the Institute for Disease Control & Prevention of the Academy of Military Medical Sciences, Beijing, China. B5233 is a clinical isolate that demonstrated a high virulence in murine infection studies, and it has been used as an experimental strain in *A. fumigatus* pathogenicity studies (35,36). The four isolates were initially identified as *A. fumigatus* by microscopic morphological description and sequencing of the internal transcribed spacer (ITS) region using Sanger sequencing.

P1MS and P1MR were originally isolated from the sputum of the same patient at different time points during a complicated Influenza A H1N1 virus infection, and were regarded as mixed infection isolates (Figure 1). This patient had no relevant underlying disease, was diagnosed with Influenza A H1N1 virus and was admitted to the UMCG. Two days after admission, a positive sputum culture of *A. fumigatus* prompted the initiation of treatment with voriconazole. Later, at day five after admission, the patient developed IA and passed away sixteen days after the diagnosis of the fungal infection. Throughout the course of the IA infection (21 days), a total of seven *A. fumigatus* isolates were recovered, where the first five isolates were susceptible to azole treatment and the last two were resistant. We selected the first susceptible and the first resistant isolates to determine their genetic relatedness.

The remaining isolate, P2CS, was recovered from an individual diagnosed with HIV and COPD. The *A. fumigatus* was cultured during a COPD exacerbation event. Chronic pulmonary aspergillosis was discarded after a chest imaging study without the radiological characteristics of pulmonary aspergillosis. Since no indicative symptoms of aspergillosis were identified, the patient was regarded as colonized by this strain. The patient is still under treatment with antiviral therapy ODEFSEY (emtricitabine/tenofovir alafenamide/rilpitvirine) and treatment for COPD with fluticason, cotrimoxazol, formeterol and ipratropium.

In addition, the raw sequencing data of five unrelated Dutch and English isolates with an environmental or clinical origin (22), were downloaded from the European Nucleotide Archive (ENA) and included in the study (Table 1).

### Antifungal susceptibility testing

To determine the *in vitro* susceptibility of the clinical isolates to triazole antifungal drugs the agar-based gradient technique for quantitative antifungal susceptibility E-test (AB BIODISK, Solna, Sweden) was used for isolates B5233 and P1MS, the agar-based method VIPcheck™ test (Nijmegen, The Netherlands) was used for isolate P2CS and the susceptibility of P1MR was determined with the in vitro EUCAST broth microdilution reference method (37).

### DNA isolation

Isolates were grown on Potato Dextrose Agar for 7 days at 35°C. DNA extraction was performed with the DNeasy UltraClean Microbioal Kit (Qiagen, Hilden, Germany) with some modifications to the initial steps of the manufacturer’s protocol. The initial fungal starting material was obtained using a pre-wetted sterile swab rubbed against the sporulating colony, that was dissolved in 700 μl sterile saline solution. The suspension was centrifuged at 10,000 rcf for 4 min. The supernatant was discarded, and the pellet was resuspended in 300 μl of Power Bead solution. This solution was used to resuspend a second pellet containing the same sample. The final concentrated solution was transferred to a microtube containing Pathogen lysis tube L beads (Qiagen, Hilden, Germany), 50 μl of Solution SL and 200 μl of sterile saline solution to homogenize. Disruption was carried out in a Tissue Homogenizer Precellys 24 (Bertin, Montigny-le-Bretonneux, France), set to 3 times 30 seconds at 5,000 rpm separated by 30 seconds. The disruption preps were heated to 65°C as suggested in the Troubleshooting Guide of the protocol to increase the final DNA yield

### Library preparation and whole genome sequencing

The whole procedure was performed according to the manufacturer’s protocol (Illumina, California, United States of America). One ng of fungal gDNA per specimen was used as input DNA for library preparation with NexteraXT DNA Library Prep Kit. Library quality was determined by measuring fragment size on a 2200 TapeStation System with D5000 Screen tape (Agilent Technologies) and quantified with Qubit 2.0 Fluorometer using Qubit dsDNA HS Assay Kit (Life Technologies, ThermoFisher Scientific, Waltham, Massachusetts, EUA). Before sequencing NexteraXT libraries were denatured and diluted to the required (equi)molarity for Illumina platform and two pools were made each containing two libraries. Whole-genome sequencing was performed in two separate runs using the MiSeq Reagent Kit v2 500-cycles Paired-End on a MiSeq Sequencer (Illumina). The raw reads generated in this study have been submitted to the European Nucleotide Archive under project accession number PRJEB28819.

### Quality control and *de-novo* assembly

The raw sequencing reads were quality trimmed using the CLC Genomics Workbench software version v10.1.1 using default settings except for the following modifications: “trim using quality scores was set to 0.01”. Assembly quality data of the nine *A. fumigatus* isolates is shown in Table S1. *De-novo* assembly produced acceptable results that surpassed a >100 coverage with >90% of reads used.

### Identification of virulence related genes

A database with genes associated with virulence was generated based on Abad *et al* review (38). In addition, these genes were validated using the online gene database AspGD (http://www.aspgd.org/). To increase the number of genes and update the database, we added secondary metabolite (SM) genes from Biosynthetic Gene Clusters (BGCs) 3, 5, 6, 14, 15 and 25, that showed a differential gene expression in murine infection studies reported by Bignell *et al* (18). A list of allergens recently revised was also included (39). A database with a total of 244 genes was created with genes categorized into seven big groups according to their site and the process they are involved in, as follows: thermotolerance, resistance to immune responses, cell wall, toxins and secondary metabolites, allergens, nutrient uptake and signaling and regulation. (Table S2) The *de-novo* assemblies of our isolates were screened with ABRicate v0.3 software tool (https://github.com/tseemann/abricate) to detect the presence or absence of VRGs included in the database. The thresholds were set to >90% coverage and >90% identity to determine the presence of a virulence gene.

### TRESP genotyping

This method is based on hypervariable Tandem Repeats located within Exons of Surface Protein coding genes (TRESP) encoding cell wall or plasma membrane proteins (20). The allele sequence repeats of three TRESP targets is combined to assign a specific genotype: an MP-2 antigenic galactomannan protein (MP2), a hypothetical protein with a CFEM domain (CFEM) and a cell surface protein A (CSP). The allele repeats of these previously described proteins were used to Create a Task Template by Allele Libraries in SeqSphere+ software v5.1.0 (Ridom GmbH, Münster, Germany) with import option: use as reference sequence “best matching allele”, which enabled a dynamic reference sequence. The assembled genomes were imported into SeqSphere+ and the specific target repetitive sequences of each protein were analyzed for each UMCG isolate using the Seqsphere+ “find in sequence” tool to identify the specific genotype.

### Comparative genomics

Genome assemblies of novel isolates were aligned using blast+ v2.6 (40) and reads were mapped with bowtie2 v2.2.5 (41) to the eight reference chromosomal genomes of Af293 (NC_007194-NC_007201). For each contig, local alignment coordinates were extended to their whole length using the highest bitscore with an in-house script. Mean coverage were calculated every 5 Kb using bedtools v2.17 (42). VRGs location were determined by local alignment and GC percentage were calculated every 100 bp with https://github.com/DamienFr/GC-content-in-sliding-window-script. Coding sequences, their location and frame where extracted from the reference sequence genbank files. All gathered information was represented in a circular image using circos v0.69-3 (43).

### Identification of variants

The variant analysis was performed for the four UMCG isolates and two Dutch environmental isolates 08-19-02-30 and 08-19-02-46. Variants were called against the reference genome *A. fumigatus* Af293 (release 37, FungiDB) using the web-based platform EuPathDB Galaxy Site(https://eupathdb.globusgenomics.org/) (44). The raw reads were quality controlled with FastQC (version 0.11.3, Brabraham institute), trimmed with Sickle (Galaxy version 070113) for quality and length thresholds of 20, aligned to reference with Bowtie2 (Align version 2.1.0 64) (45) using the “very sensitive” alignment preset option, BAM file sorted with SAMtools and variant calling performed with Freebayes (v0.9.21-19-gc003c1e) and SAMtools (46). The resulting variants were annotated using SnpEff to predict the impact of a variant on the effect of a gene, by classifying them in different categories: high, moderate, low and modifier (47). (http://snpeff.sourceforge.net/SnpEff_manual.html)High impact variants are predicted to have a disruptive effect in the protein (e.g. frameshift variants, inversion), moderate impact variants could change the protein effectiveness (e.g. missense variant, in frame deletion), low impact variants are not expected to have a big impact in the protein function (e.g. synonymous variant) and finally, modifier variants are non-coding changes where predictions are difficult or there is no evidence of impact (e.g. exon variant, downstream gene variant). SnpSift was used to extract the variants with moderate and high impact by filtering the resulting variant call format (VCF) files from SnpEeff.

In addition, identification of SNPs present in VRGs of our isolates, was performed using CLC Genomics Workbench software version v11.0.1. For this approach, trimmed reads of each genome were mapped to a concatenated sequence consisted of 244 VRG genes described in Table S2. SNPs were called with a minimum read coverage of 10 and with a minimum frequency of 90%. The used virulence gene sequences to create the concatenated sequence belonged to the reference *A. fumigatus* Af293.

Snippy v. 4.3.5 (https://github.com/tseemann/snippy) was used to determine the number of variants between isolate P1MS and P1MR. The trimmed reads of isolate P1MR were aligned to the assembly of P1MS for variant calling. In this case, P1MS draft genome assembly, used as reference, is not annotated and therefore, a functional prediction of the determined variants was not possible and only a quantitative analysis is presented.

## Acknowledgements

F.P.B was supported by the Erasmus Mundus Joint Master Degree (EMJMD) scholarship of the Erasmus+ EU-Programme awarded under the International Master in Innovative Medicine (IMIM) programme. We thank Xuelin Han and Li Han from the Institute for Disease Control & Prevention of the Academy of Military Medical Sciences, Beijing, China for providing strain B5233 and K. J. Kwon-Chung for providing information about the origin of this isolate.

This work was partly supported by the INTERREG VA (202085) funded project EurHealth-1Health, part of a Dutch-German cross-border network supported by the European Commission, the Dutch Ministry of Health, Welfare and Sport (VWS), the Ministry of Economy, Innovation, Digitalisation and Energy of the German Federal State of North Rhine-Westphalia and the German Federal State of Lower Saxony.

---

**TABLE S1.**
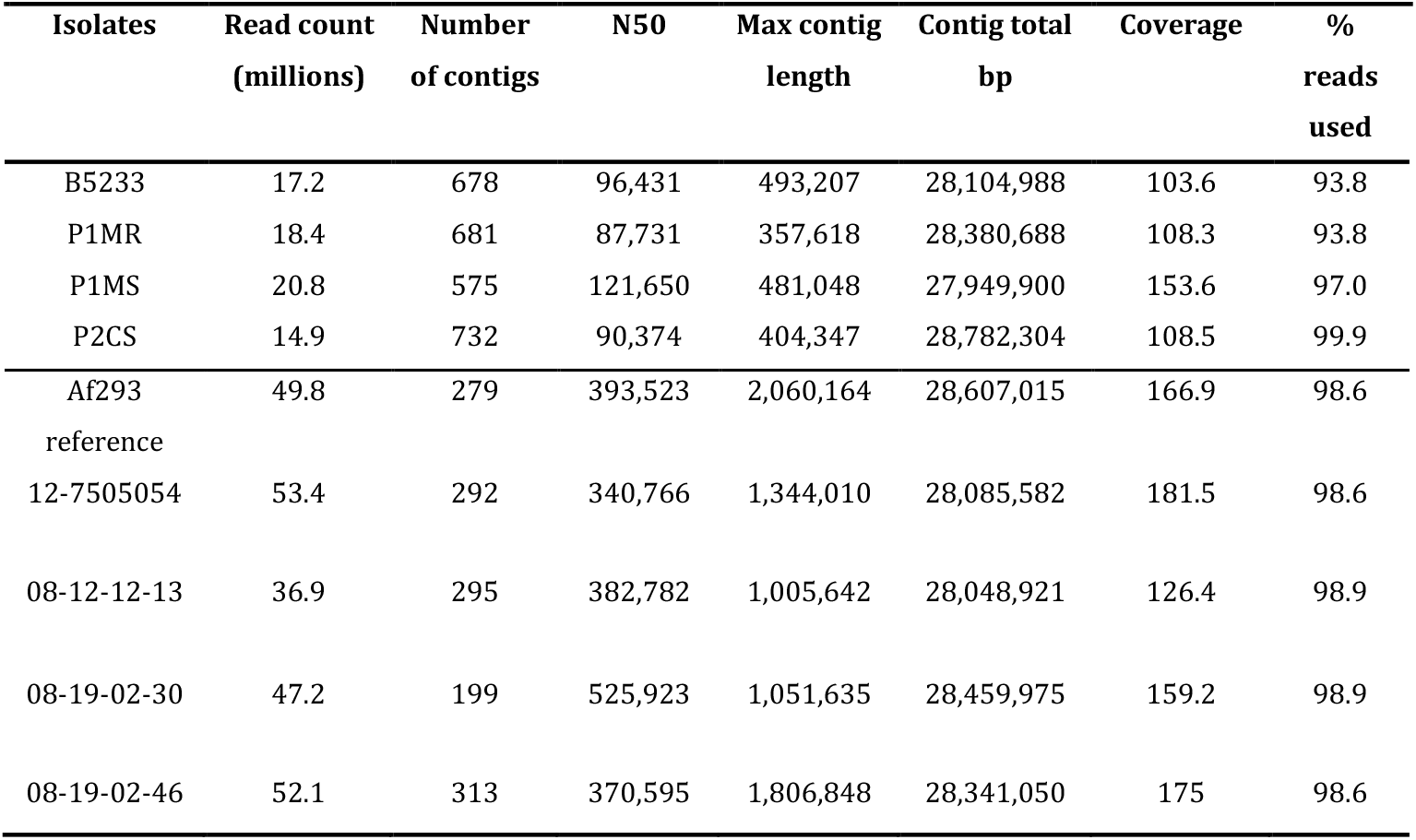
Summary of *de-novo* assemblies using CLC Workbench v10.1.1.

**TABLE S2.**
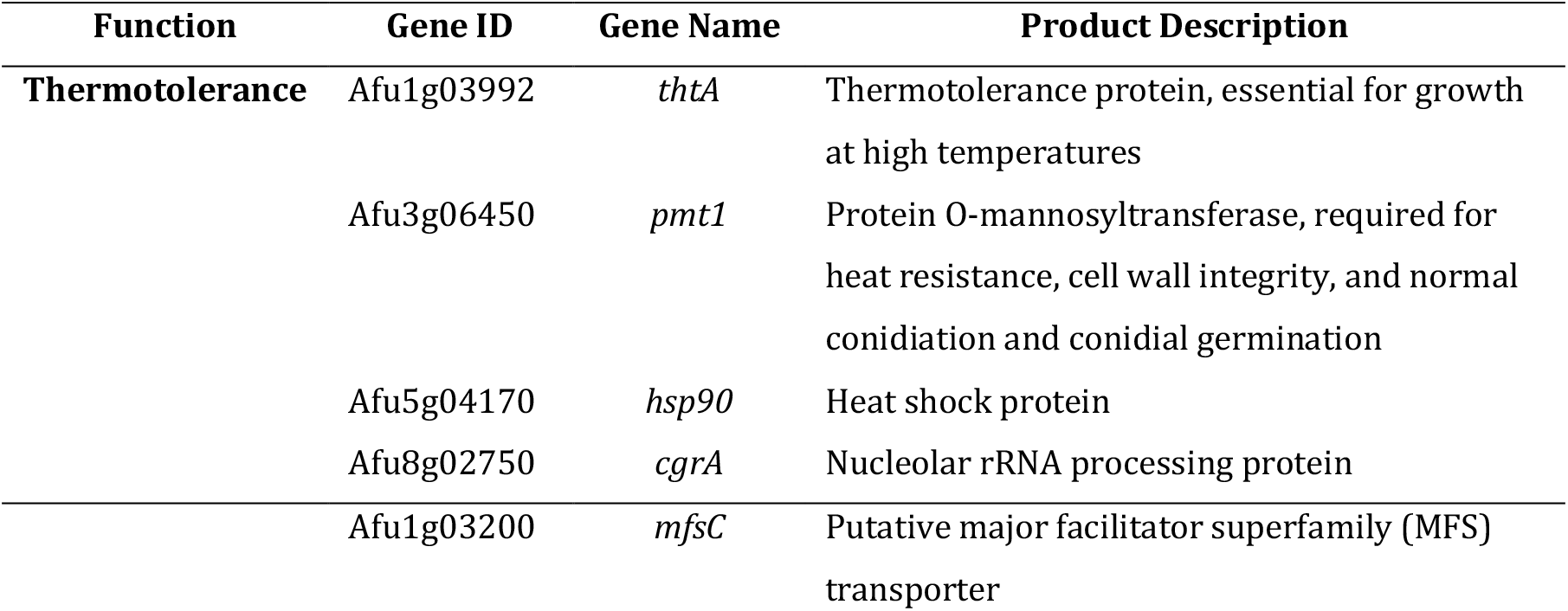
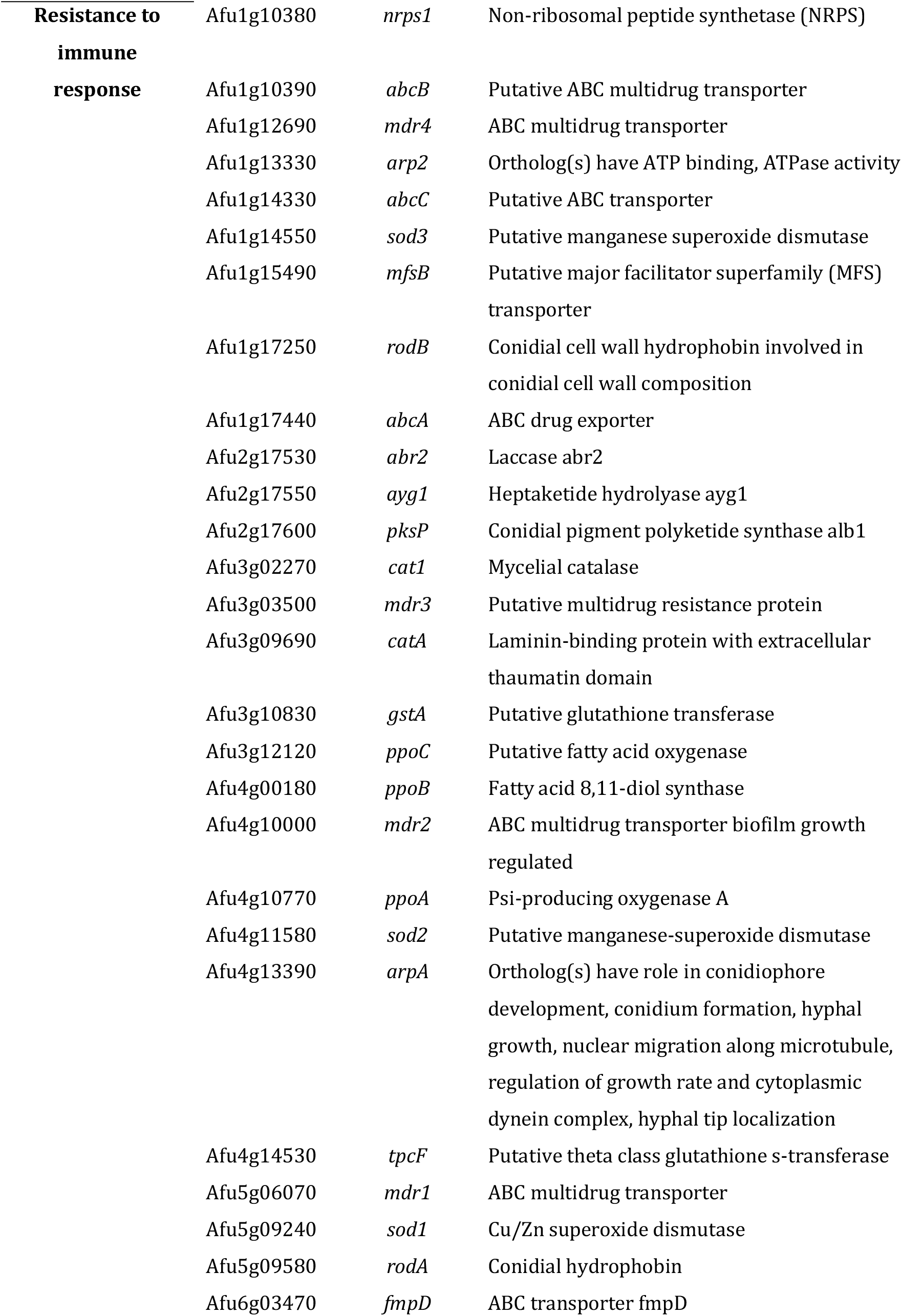
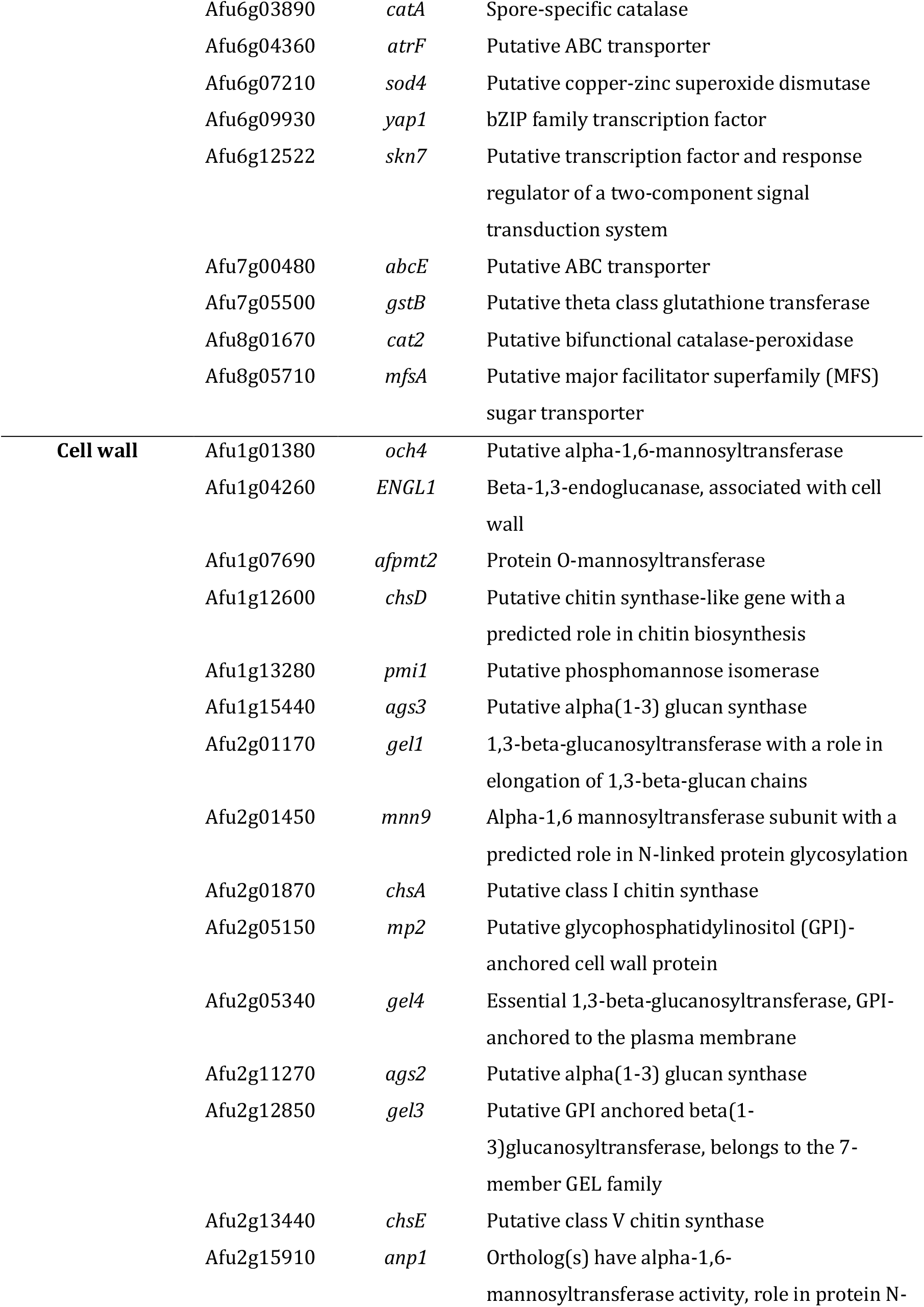
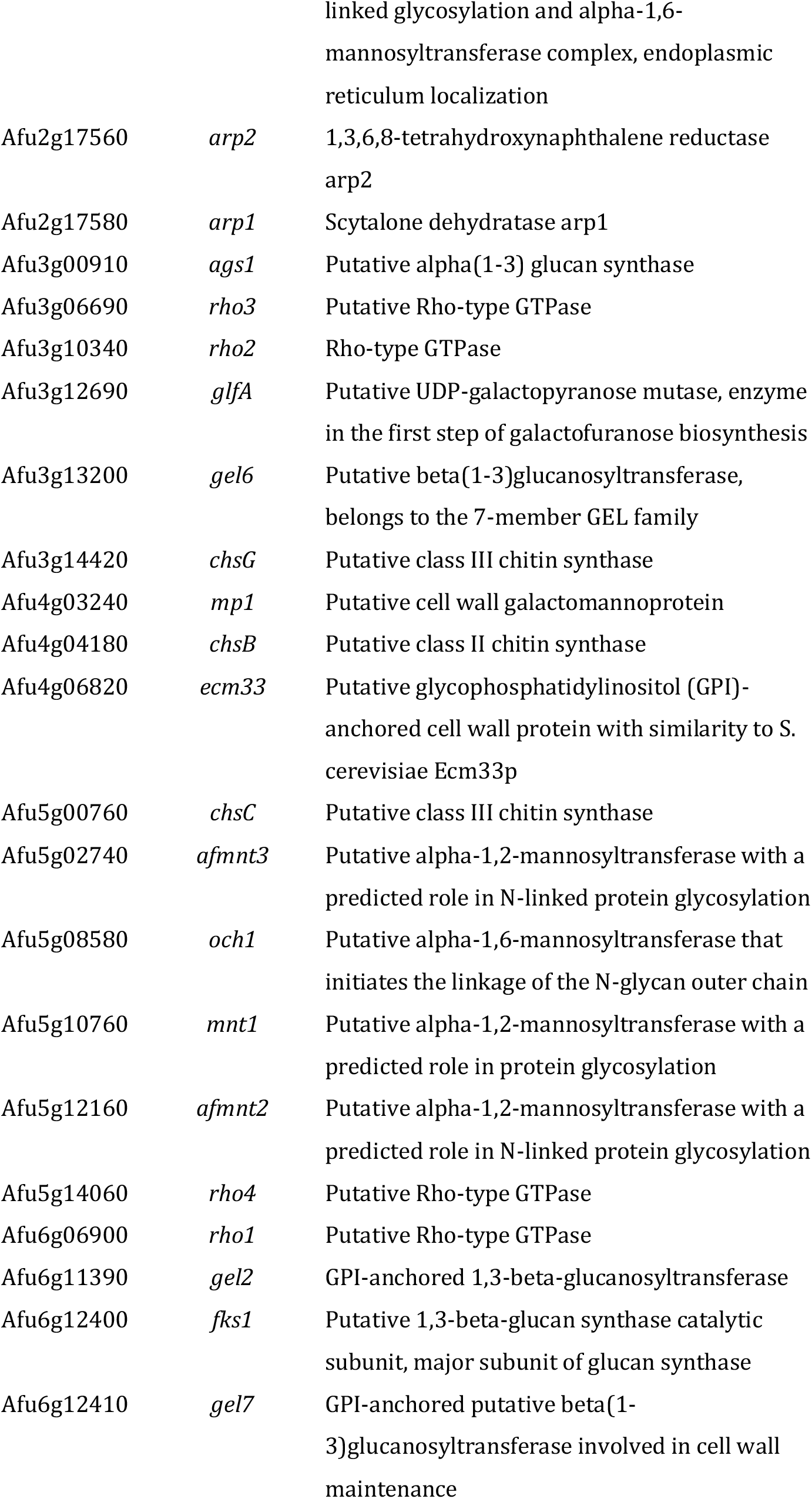
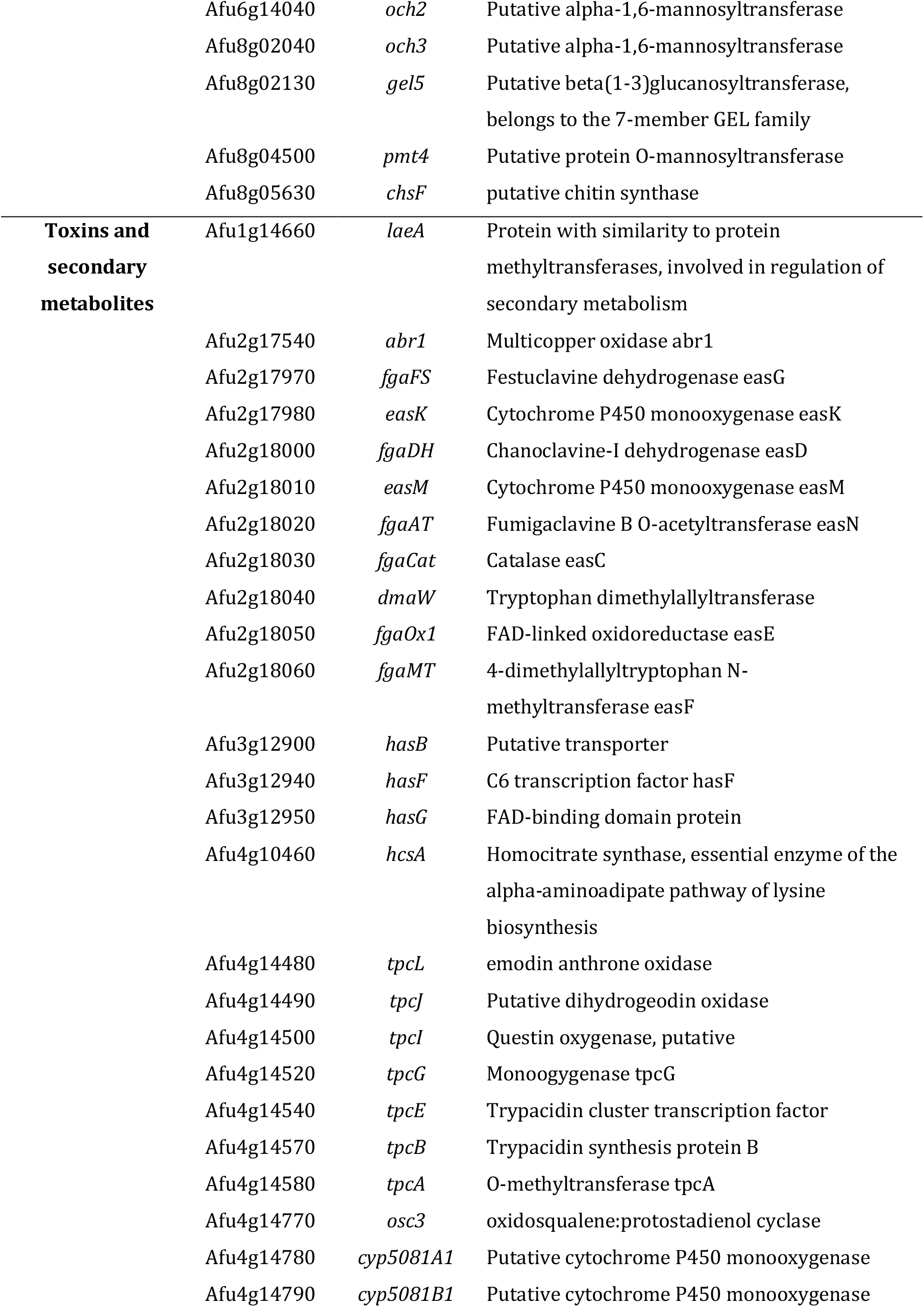
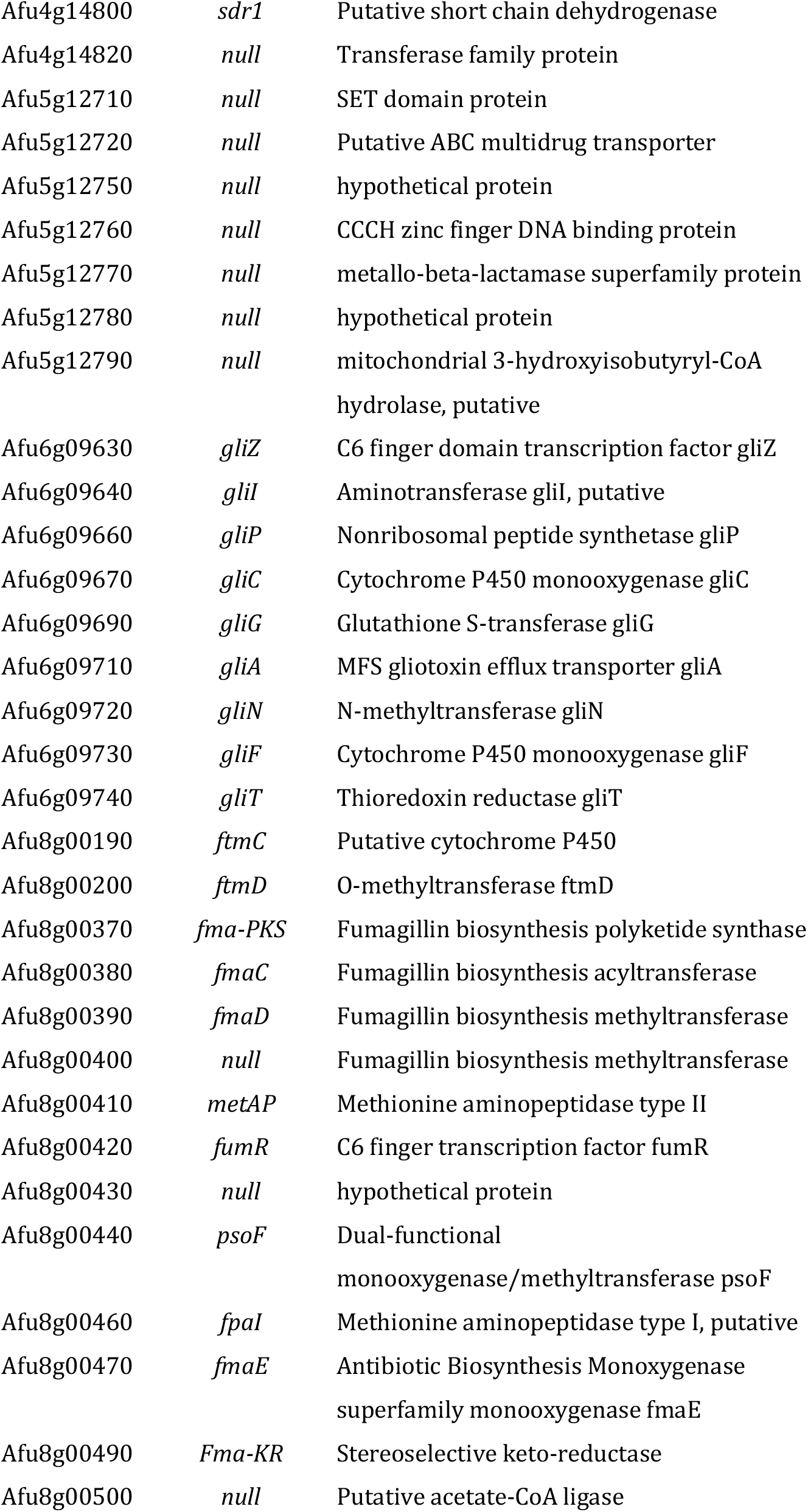
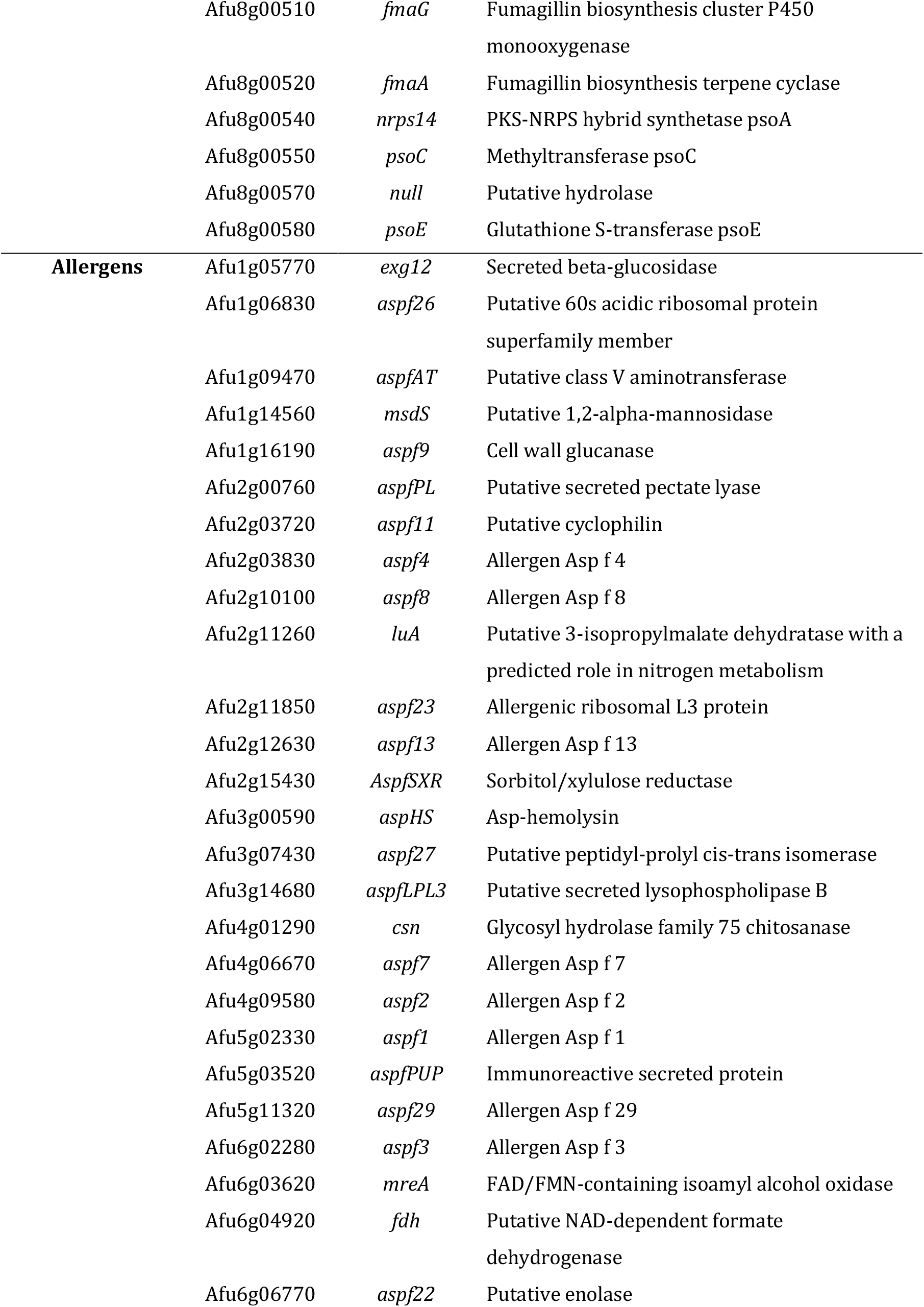
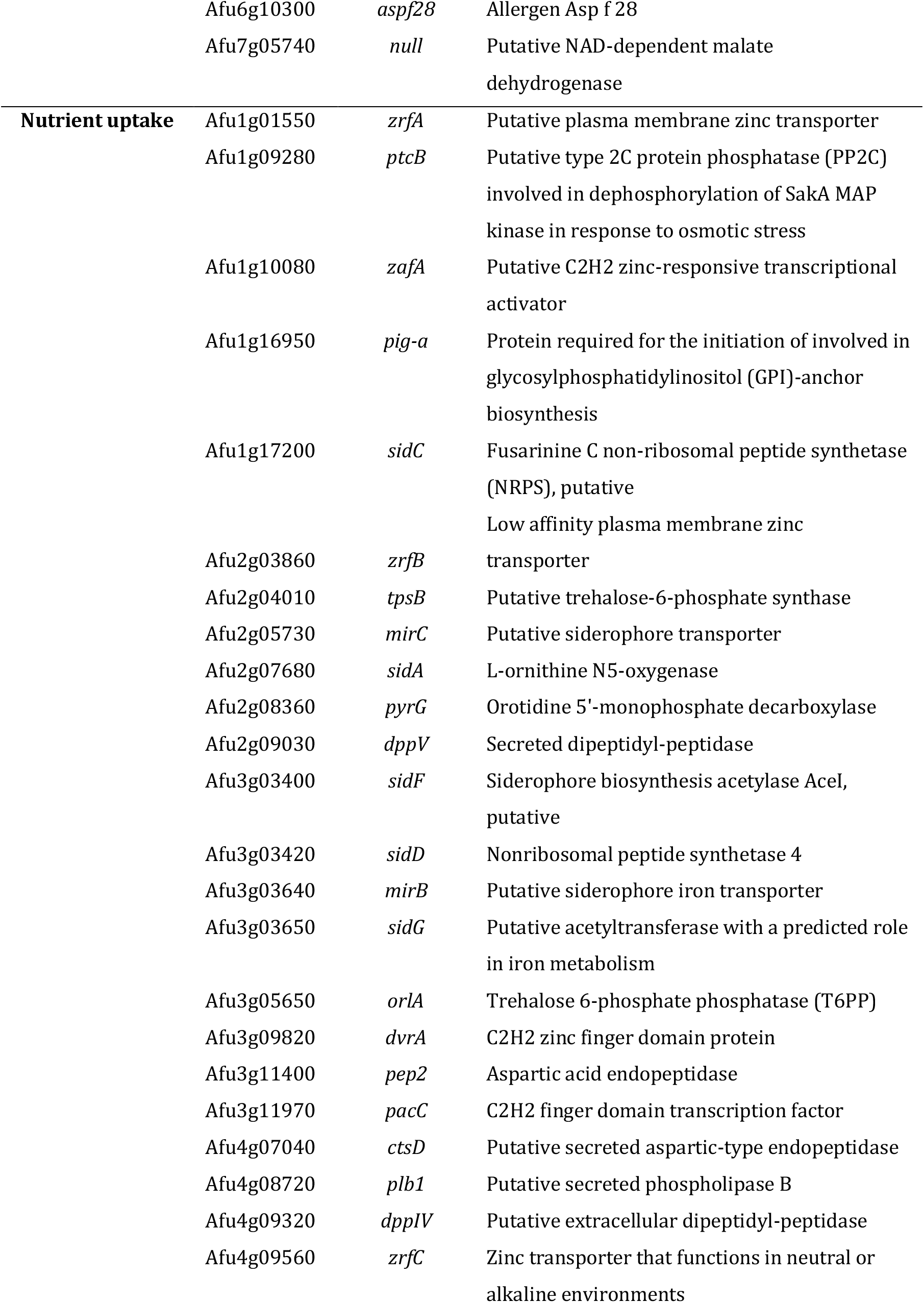
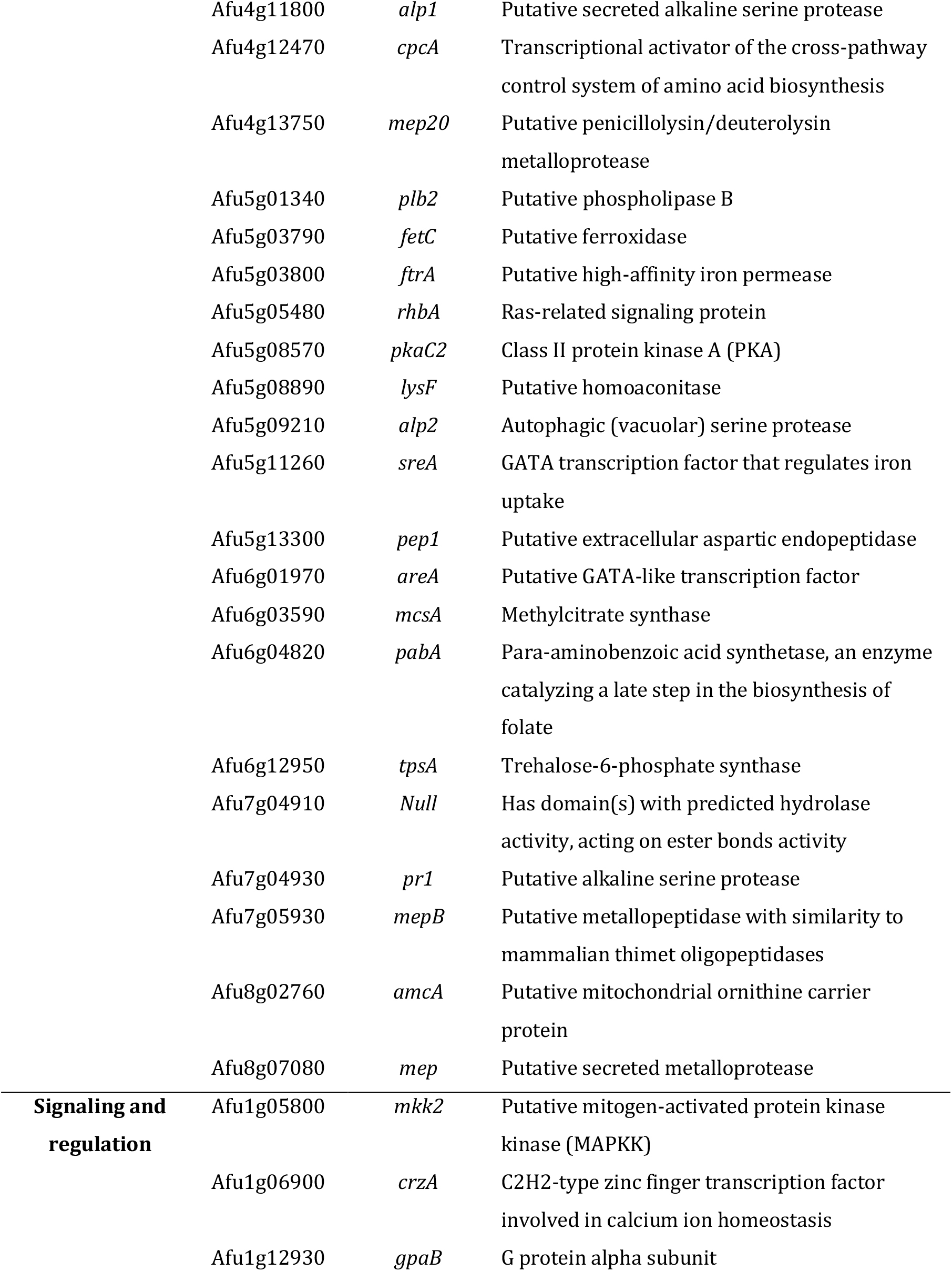
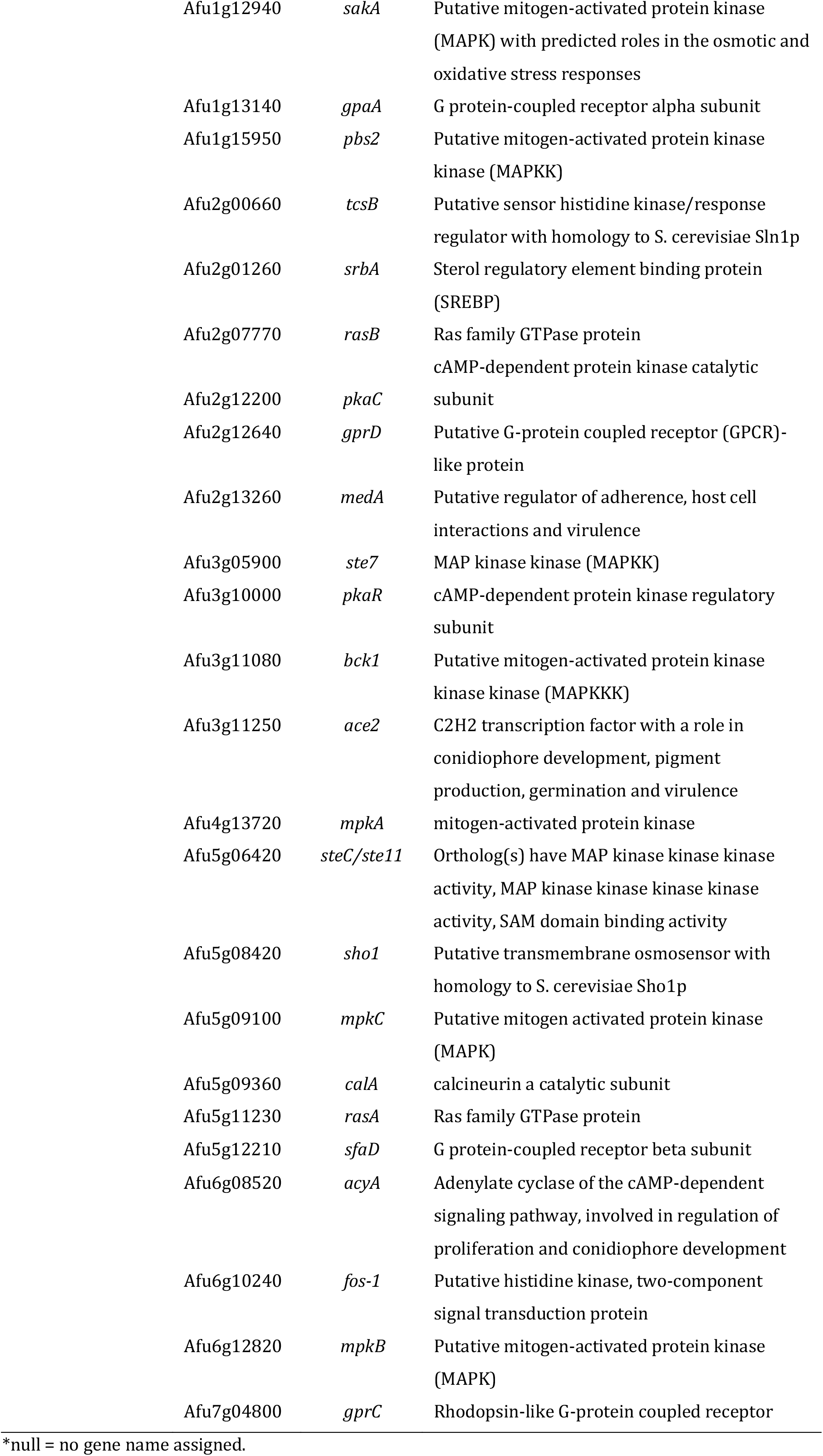
Virulence related genes included in our in-house database for the screening of *A. fumigatus* isolates.

**Table S3.**
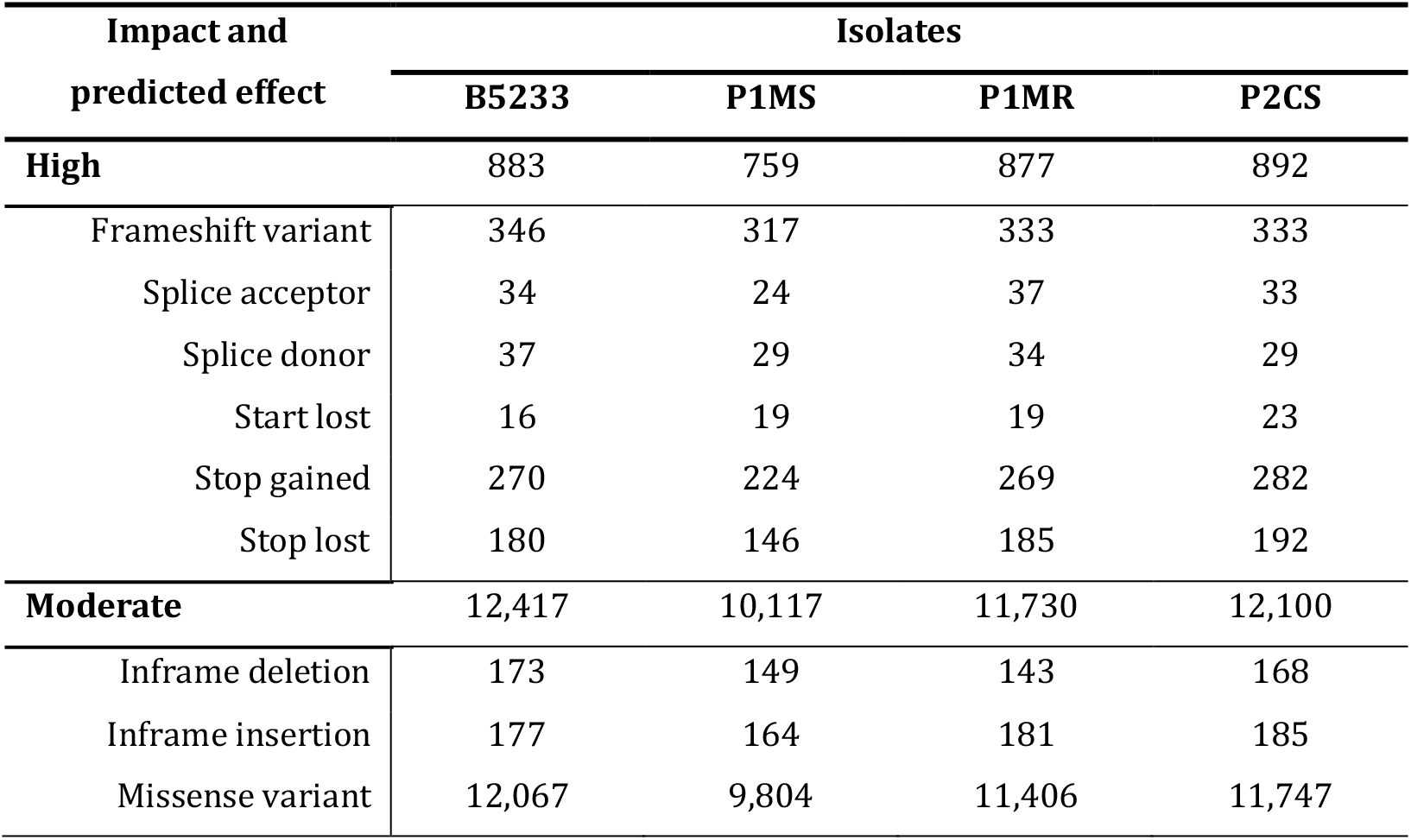
Predicted effect of resulting variants after filtering VCF files with SnpSift.

