## Supplementary material for "Revealing virulence potential of clinical and environmental *Aspergillus fumigatus* isolates using Whole-genome sequencing"

**TABLE S1.** Summary of d*e-novo* assemblies using CLC Workbench v10.1.1.

| Isolates | Read count (millions) | Number of contigs | N50 | Max contig length | Contig total bp | Coverage | % reads used |
|---|---|---|---|---|---|---|---|
| B5233 | 17.2 | 678 | 96,431 | 493,207 | 28,104,988 | 103.6 | 93.8 |
| P1MR | 18.4 | 681 | 87,731 | 357,618 | 28,380,688 | 108.3 | 93.8 |
| P1MS | 20.8 | 575 | 121,650 | 481,048 | 27,949,900 | 153.6 | 97.0 |
| P2CS | 14.9 | 732 | 90,374 | 404,347 | 28,782,304 | 108.5 | 99.9 |
| Af293 reference | 49.8 | 279 | 393,523 | 2,060,164 | 28,607,015 | 166.9 | 98.6 |
| 12-7505054 | 53.4 | 292 | 340,766 | 1,344,010 | 28,085,582 | 181.5 | 98.6 |
| 08-12-12-13 | 36.9 | 295 | 382,782 | 1,005,642 | 28,048,921 | 126.4 | 98.9 |
| 08-19-02-30 | 47.2 | 199 | 525,923 | 1,051,635 | 28,459,975 | 159.2 | 98.9 |
| 08-19-02-46 | 52.1 | 313 | 370,595 | 1,806,848 | 28,341,050 | 175 | 98.6 |

**TABLE S2.** Virulence related genes included in our in-house database for the screening of *A. fumigatus* isolates.

| Function | Gene ID | Gene Name | Product Description |
|---|---|---|---|
| **Thermotolerance** | Afu1g03992 | *thtA* | Thermotolerance protein, essential for growth at high temperatures |
| | Afu3g06450 | *pmt1* | Protein O-mannosyltransferase, required for heat resistance, cell wall integrity, and normal conidiation and conidial germination |
| | Afu5g04170 | *hsp90* | Heat shock protein |
| | Afu8g02750 | *cgrA* | Nucleolar rRNA processing protein |
| **Resistance to immune response** | Afu1g03200 | *mfsC* | Putative major facilitator superfamily (MFS) transporter |
| | Afu1g10380 | *nrps1* | Non-ribosomal peptide synthetase (NRPS) |
| | Afu1g10390 | *abcB* | Putative ABC multidrug transporter |
| | Afu1g12690 | *mdr4* | ABC multidrug transporter |
| | Afu1g13330 | *arp2* | Ortholog(s) have ATP binding, ATPase activity |
| | Afu1g14330 | *abcC* | Putative ABC transporter |
| | Afu1g14550 | *sod3* | Putative manganese superoxide dismutase |
| | Afu1g15490 | *mfsB* | Putative major facilitator superfamily (MFS) transporter |

| | | |
|---|---|---|
| Afu1g17250 | *rodB* | Conidial cell wall hydrophobin involved in conidial cell wall composition |
| Afu1g17440 | *abcA* | ABC drug exporter |
| Afu2g17530 | *abr2* | Laccase abr2 |
| Afu2g17550 | *ayg1* | Heptaketide hydrolyase ayg1 |
| Afu2g17600 | *pksP* | Conidial pigment polyketide synthase alb1 |
| Afu3g02270 | *cat1* | Mycelial catalase |
| Afu3g03500 | *mdr3* | Putative multidrug resistance protein |
| Afu3g09690 | *catA* | Laminin-binding protein with extracellular thaumatin domain |
| Afu3g10830 | *gstA* | Putative glutathione transferase |
| Afu3g12120 | *ppoC* | Putative fatty acid oxygenase |
| Afu4g00180 | *ppoB* | Fatty acid 8,11-diol synthase |
| Afu4g10000 | *mdr2* | ABC multidrug transporter biofilm growth regulated |
| Afu4g10770 | *ppoA* | Psi-producing oxygenase A |
| Afu4g11580 | *sod2* | Putative manganese-superoxide dismutase |
| Afu4g13390 | *arpA* | Ortholog(s) have role in conidiophore development, conidium formation, hyphal growth, nuclear migration along microtubule, regulation of growth rate and cytoplasmic dynein complex, hyphal tip localization |
| Afu4g14530 | *tpcF* | Putative theta class glutathione s-transferase |
| Afu5g06070 | *mdr1* | ABC multidrug transporter |
| Afu5g09240 | *sod1* | Cu/Zn superoxide dismutase |
| Afu5g09580 | *rodA* | Conidial hydrophobin |
| Afu6g03470 | *fmpD* | ABC transporter fmpD |
| Afu6g03890 | *catA* | Spore-specific catalase |
| Afu6g04360 | *atrF* | Putative ABC transporter |
| Afu6g07210 | *sod4* | Putative copper-zinc superoxide dismutase |
| Afu6g09930 | *yap1* | bZIP family transcription factor |
| Afu6g12522 | *skn7* | Putative transcription factor and response regulator of a two-component signal transduction system |
| Afu7g00480 | *abcE* | Putative ABC transporter |
| Afu7g05500 | *gstB* | Putative theta class glutathione transferase |
| Afu8g01670 | *cat2* | Putative bifunctional catalase-peroxidase |
| Afu8g05710 | *mfsA* | Putative major facilitator superfamily (MFS) sugar transporter |

| | | | |
|---|---|---|---|
| **Cell wall** | Afu1g01380 | *och4* | Putative alpha-1,6-mannosyltransferase |
| | Afu1g04260 | *ENGL1* | Beta-1,3-endoglucanase, associated with cell wall |
| | Afu1g07690 | *afpmt2* | Protein O-mannosyltransferase |
| | Afu1g12600 | *chsD* | Putative chitin synthase-like gene with a predicted role in chitin biosynthesis |
| | Afu1g13280 | *pmi1* | Putative phosphomannose isomerase |
| | Afu1g15440 | *ags3* | Putative alpha(1-3) glucan synthase |
| | Afu2g01170 | *gel1* | 1,3-beta-glucanosyltransferase with a role in elongation of 1,3-beta-glucan chains |
| | Afu2g01450 | *mnn9* | Alpha-1,6 mannosyltransferase subunit with a predicted role in N-linked protein glycosylation |
| | Afu2g01870 | *chsA* | Putative class I chitin synthase |
| | Afu2g05150 | *mp2* | Putative glycophosphatidylinositol (GPI)-anchored cell wall protein |
| | Afu2g05340 | *gel4* | Essential 1,3-beta-glucanosyltransferase, GPI-anchored to the plasma membrane |
| | Afu2g11270 | *ags2* | Putative alpha(1-3) glucan synthase |
| | Afu2g12850 | *gel3* | Putative GPI anchored beta(1-3)glucanosyltransferase, belongs to the 7-member GEL family |
| | Afu2g13440 | *chsE* | Putative class V chitin synthase |
| | Afu2g15910 | *anp1* | Ortholog(s) have alpha-1,6-mannosyltransferase activity, role in protein N-linked glycosylation and alpha-1,6-mannosyltransferase complex, endoplasmic reticulum localization |
| | Afu2g17560 | *arp2* | 1,3,6,8-tetrahydroxynaphthalene reductase arp2 |
| | Afu2g17580 | *arp1* | Scytalone dehydratase arp1 |
| | Afu3g00910 | *ags1* | Putative alpha(1-3) glucan synthase |
| | Afu3g06690 | *rho3* | Putative Rho-type GTPase |
| | Afu3g10340 | *rho2* | Rho-type GTPase |
| | Afu3g12690 | *glfA* | Putative UDP-galactopyranose mutase, enzyme in the first step of galactofuranose biosynthesis |
| | Afu3g13200 | *gel6* | Putative beta(1-3)glucanosyltransferase, belongs to the 7-member GEL family |
| | Afu3g14420 | *chsG* | Putative class III chitin synthase |
| | Afu4g03240 | *mp1* | Putative cell wall galactomannoprotein |

| | | | |
|---|---|---|---|
| | Afu4g04180 | *chsB* | Putative class II chitin synthase |
| | Afu4g06820 | *ecm33* | Putative glycophosphatidylinositol (GPI)-anchored cell wall protein with similarity to S. cerevisiae Ecm33p |
| | Afu5g00760 | *chsC* | Putative class III chitin synthase |
| | Afu5g02740 | *afmnt3* | Putative alpha-1,2-mannosyltransferase with a predicted role in N-linked protein glycosylation |
| | Afu5g08580 | *och1* | Putative alpha-1,6-mannosyltransferase that initiates the linkage of the N-glycan outer chain |
| | Afu5g10760 | *mnt1* | Putative alpha-1,2-mannosyltransferase with a predicted role in protein glycosylation |
| | Afu5g12160 | *afmnt2* | Putative alpha-1,2-mannosyltransferase with a predicted role in N-linked protein glycosylation |
| | Afu5g14060 | *rho4* | Putative Rho-type GTPase |
| | Afu6g06900 | *rho1* | Putative Rho-type GTPase |
| | Afu6g11390 | *gel2* | GPI-anchored 1,3-beta-glucanosyltransferase |
| | Afu6g12400 | *fks1* | Putative 1,3-beta-glucan synthase catalytic subunit, major subunit of glucan synthase |
| | Afu6g12410 | *gel7* | GPI-anchored putative beta(1-3)glucanosyltransferase involved in cell wall maintenance |
| | Afu6g14040 | *och2* | Putative alpha-1,6-mannosyltransferase |
| | Afu8g02040 | *och3* | Putative alpha-1,6-mannosyltransferase |
| | Afu8g02130 | *gel5* | Putative beta(1-3)glucanosyltransferase, belongs to the 7-member GEL family |
| | Afu8g04500 | *pmt4* | Putative protein O-mannosyltransferase |
| | Afu8g05630 | *chsF* | putative chitin synthase |
| **Toxins and secondary metabolites** | Afu1g14660 | *laeA* | Protein with similarity to protein methyltransferases, involved in regulation of secondary metabolism |
| | Afu2g17540 | *abr1* | Multicopper oxidase abr1 |
| | Afu2g17970 | *fgaFS* | Festuclavine dehydrogenase easG |
| | Afu2g17980 | *easK* | Cytochrome P450 monooxygenase easK |
| | Afu2g18000 | *fgaDH* | Chanoclavine-I dehydrogenase easD |
| | Afu2g18010 | *easM* | Cytochrome P450 monooxygenase easM |
| | Afu2g18020 | *fgaAT* | Fumigaclavine B O-acetyltransferase easN |
| | Afu2g18030 | *fgaCat* | Catalase easC |
| | Afu2g18040 | *dmaW* | Tryptophan dimethylallyltransferase |
| | Afu2g18050 | *fgaOx1* | FAD-linked oxidoreductase easE |

| | | |
|---|---|---|
| Afu2g18060 | *fgaMT* | 4-dimethylallyltryptophan N-methyltransferase easF |
| Afu3g12900 | *hasB* | Putative transporter |
| Afu3g12940 | *hasF* | C6 transcription factor hasF |
| Afu3g12950 | *hasG* | FAD-binding domain protein |
| Afu4g10460 | *hcsA* | Homocitrate synthase, essential enzyme of the alpha-aminoadipate pathway of lysine biosynthesis |
| Afu4g14480 | *tpcL* | emodin anthrone oxidase |
| Afu4g14490 | *tpcJ* | Putative dihydrogeodin oxidase |
| Afu4g14500 | *tpcI* | Questin oxygenase, putative |
| Afu4g14520 | *tpcG* | Monoogygenase tpcG |
| Afu4g14540 | *tpcE* | Trypacidin cluster transcription factor |
| Afu4g14570 | *tpcB* | Trypacidin synthesis protein B |
| Afu4g14580 | *tpcA* | O-methyltransferase tpcA |
| Afu4g14770 | *osc3* | oxidosqualene:protostadienol cyclase |
| Afu4g14780 | *cyp5081A1* | Putative cytochrome P450 monooxygenase |
| Afu4g14790 | *cyp5081B1* | Putative cytochrome P450 monooxygenase |
| Afu4g14800 | *sdr1* | Putative short chain dehydrogenase |
| Afu4g14820 | *null* | Transferase family protein |
| Afu5g12710 | *null* | SET domain protein |
| Afu5g12720 | *null* | Putative ABC multidrug transporter |
| Afu5g12750 | *null* | hypothetical protein |
| Afu5g12760 | *null* | CCCH zinc finger DNA binding protein |
| Afu5g12770 | *null* | metallo-beta-lactamase superfamily protein |
| Afu5g12780 | *null* | hypothetical protein |
| Afu5g12790 | *null* | mitochondrial 3-hydroxyisobutyryl-CoA hydrolase, putative |
| Afu6g09630 | *gliZ* | C6 finger domain transcription factor gliZ |
| Afu6g09640 | *gliI* | Aminotransferase gliI, putative |
| Afu6g09660 | *gliP* | Nonribosomal peptide synthetase gliP |
| Afu6g09670 | *gliC* | Cytochrome P450 monooxygenase gliC |
| Afu6g09690 | *gliG* | Glutathione S-transferase gliG |
| Afu6g09710 | *gliA* | MFS gliotoxin efflux transporter gliA |
| Afu6g09720 | *gliN* | N-methyltransferase gliN |
| Afu6g09730 | *gliF* | Cytochrome P450 monooxygenase gliF |
| Afu6g09740 | *gliT* | Thioredoxin reductase gliT |
| Afu8g00190 | *ftmC* | Putative cytochrome P450 |
| Afu8g00200 | *ftmD* | O-methyltransferase ftmD |

| | | | |
|---|---|---|---|
| | Afu8g00370 | *fma-PKS* | Fumagillin biosynthesis polyketide synthase |
| | Afu8g00380 | *fmaC* | Fumagillin biosynthesis acyltransferase |
| | Afu8g00390 | *fmaD* | Fumagillin biosynthesis methyltransferase |
| | Afu8g00400 | *null* | Fumagillin biosynthesis methyltransferase |
| | Afu8g00410 | *metAP* | Methionine aminopeptidase type II |
| | Afu8g00420 | *fumR* | C6 finger transcription factor fumR |
| | Afu8g00430 | *null* | hypothetical protein |
| | Afu8g00440 | *psoF* | Dual-functional monooxygenase/methyltransferase psoF |
| | Afu8g00460 | *fpaI* | Methionine aminopeptidase type I, putative |
| | Afu8g00470 | *fmaE* | Antibiotic Biosynthesis Monoxygenase superfamily monooxygenase fmaE |
| | Afu8g00490 | *Fma-KR* | Stereoselective keto-reductase |
| | Afu8g00500 | *null* | Putative acetate-CoA ligase |
| | Afu8g00510 | *fmaG* | Fumagillin biosynthesis cluster P450 monooxygenase |
| | Afu8g00520 | *fmaA* | Fumagillin biosynthesis terpene cyclase |
| | Afu8g00540 | *nrps14* | PKS-NRPS hybrid synthetase psoA |
| | Afu8g00550 | *psoC* | Methyltransferase psoC |
| | Afu8g00570 | *null* | Putative hydrolase |
| | Afu8g00580 | *psoE* | Glutathione S-transferase psoE |
| **Allergens** | Afu1g05770 | *exg12* | Secreted beta-glucosidase |
| | Afu1g06830 | *aspf26* | Putative 60s acidic ribosomal protein superfamily member |
| | Afu1g09470 | *aspfAT* | Putative class V aminotransferase |
| | Afu1g14560 | *msdS* | Putative 1,2-alpha-mannosidase |
| | Afu1g16190 | *aspf9* | Cell wall glucanase |
| | Afu2g00760 | *aspfPL* | Putative secreted pectate lyase |
| | Afu2g03720 | *aspf11* | Putative cyclophilin |
| | Afu2g03830 | *aspf4* | Allergen Asp f 4 |
| | Afu2g10100 | *aspf8* | Allergen Asp f 8 |
| | Afu2g11260 | *luA* | Putative 3-isopropylmalate dehydratase with a predicted role in nitrogen metabolism |
| | Afu2g11850 | *aspf23* | Allergenic ribosomal L3 protein |
| | Afu2g12630 | *aspf13* | Allergen Asp f 13 |
| | Afu2g15430 | *AspfSXR* | Sorbitol/xylulose reductase |
| | Afu3g00590 | *aspHS* | Asp-hemolysin |
| | Afu3g07430 | *aspf27* | Putative peptidyl-prolyl cis-trans isomerase |
| | Afu3g14680 | *aspfLPL3* | Putative secreted lysophospholipase B |

| | | | |
|---|---|---|---|
| | Afu4g01290 | *csn* | Glycosyl hydrolase family 75 chitosanase |
| | Afu4g06670 | *aspf7* | Allergen Asp f 7 |
| | Afu4g09580 | *aspf2* | Allergen Asp f 2 |
| | Afu5g02330 | *aspf1* | Allergen Asp f 1 |
| | Afu5g03520 | *aspfPUP* | Immunoreactive secreted protein |
| | Afu5g11320 | *aspf29* | Allergen Asp f 29 |
| | Afu6g02280 | *aspf3* | Allergen Asp f 3 |
| | Afu6g03620 | *mreA* | FAD/FMN-containing isoamyl alcohol oxidase |
| | Afu6g04920 | *fdh* | Putative NAD-dependent formate dehydrogenase |
| | Afu6g06770 | *aspf22* | Putative enolase |
| | Afu6g10300 | *aspf28* | Allergen Asp f 28 |
| | Afu7g05740 | *null* | Putative NAD-dependent malate dehydrogenase |
| **Nutrient uptake** | Afu1g01550 | *zrfA* | Putative plasma membrane zinc transporter |
| | Afu1g09280 | *ptcB* | Putative type 2C protein phosphatase (PP2C) involved in dephosphorylation of SakA MAP kinase in response to osmotic stress |
| | Afu1g10080 | *zafA* | Putative C2H2 zinc-responsive transcriptional activator |
| | Afu1g16950 | *pig-a* | Protein required for the initiation of involved in glycosylphosphatidylinositol (GPI)-anchor biosynthesis |
| | Afu1g17200 | *sidC* | Fusarinine C non-ribosomal peptide synthetase (NRPS), putative |
| | Afu2g03860 | *zrfB* | Low affinity plasma membrane zinc transporter |
| | Afu2g04010 | *tpsB* | Putative trehalose-6-phosphate synthase |
| | Afu2g05730 | *mirC* | Putative siderophore transporter |
| | Afu2g07680 | *sidA* | L-ornithine N5-oxygenase |
| | Afu2g08360 | *pyrG* | Orotidine 5'-monophosphate decarboxylase |
| | Afu2g09030 | *dppV* | Secreted dipeptidyl-peptidase |
| | Afu3g03400 | *sidF* | Siderophore biosynthesis acetylase AceI, putative |
| | Afu3g03420 | *sidD* | Nonribosomal peptide synthetase 4 |
| | Afu3g03640 | *mirB* | Putative siderophore iron transporter |
| | Afu3g03650 | *sidG* | Putative acetyltransferase with a predicted role in iron metabolism |
| | Afu3g05650 | *orlA* | Trehalose 6-phosphate phosphatase (T6PP) |
| | Afu3g09820 | *dvrA* | C2H2 zinc finger domain protein |

| | | | |
|---|---|---|---|
| | Afu3g11400 | *pep2* | Aspartic acid endopeptidase |
| | Afu3g11970 | *pacC* | C2H2 finger domain transcription factor |
| | Afu4g07040 | *ctsD* | Putative secreted aspartic-type endopeptidase |
| | Afu4g08720 | *plb1* | Putative secreted phospholipase B |
| | Afu4g09320 | *dppIV* | Putative extracellular dipeptidyl-peptidase |
| | Afu4g09560 | *zrfC* | Zinc transporter that functions in neutral or alkaline environments |
| | Afu4g11800 | *alp1* | Putative secreted alkaline serine protease |
| | Afu4g12470 | *cpcA* | Transcriptional activator of the cross-pathway control system of amino acid biosynthesis |
| | Afu4g13750 | *mep20* | Putative penicillolysin/deuterolysin metalloprotease |
| | Afu5g01340 | *plb2* | Putative phospholipase B |
| | Afu5g03790 | *fetC* | Putative ferroxidase |
| | Afu5g03800 | *ftrA* | Putative high-affinity iron permease |
| | Afu5g05480 | *rhbA* | Ras-related signaling protein |
| | Afu5g08570 | *pkaC2* | Class II protein kinase A (PKA) |
| | Afu5g08890 | *lysF* | Putative homoaconitase |
| | Afu5g09210 | *alp2* | Autophagic (vacuolar) serine protease |
| | Afu5g11260 | *sreA* | GATA transcription factor that regulates iron uptake |
| | Afu5g13300 | *pep1* | Putative extracellular aspartic endopeptidase |
| | Afu6g01970 | *areA* | Putative GATA-like transcription factor |
| | Afu6g03590 | *mcsA* | Methylcitrate synthase |
| | Afu6g04820 | *pabA* | Para-aminobenzoic acid synthetase, an enzyme catalyzing a late step in the biosynthesis of folate |
| | Afu6g12950 | *tpsA* | Trehalose-6-phosphate synthase |
| | Afu7g04910 | *Null* | Has domain(s) with predicted hydrolase activity, acting on ester bonds activity |
| | Afu7g04930 | *pr1* | Putative alkaline serine protease |
| | Afu7g05930 | *mepB* | Putative metallopeptidase with similarity to mammalian thimet oligopeptidases |
| | Afu8g02760 | *amcA* | Putative mitochondrial ornithine carrier protein |
| | Afu8g07080 | *mep* | Putative secreted metalloprotease |
| **Signaling and regulation** | Afu1g05800 | *mkk2* | Putative mitogen-activated protein kinase kinase (MAPKK) |

| | | |
|---|---|---|
| Afu1g06900 | *crzA* | C2H2-type zinc finger transcription factor involved in calcium ion homeostasis |
| Afu1g12930 | *gpaB* | G protein alpha subunit |
| Afu1g12940 | *sakA* | Putative mitogen-activated protein kinase (MAPK) with predicted roles in the osmotic and oxidative stress responses |
| Afu1g13140 | *gpaA* | G protein-coupled receptor alpha subunit |
| Afu1g15950 | *pbs2* | Putative mitogen-activated protein kinase kinase (MAPKK) |
| Afu2g00660 | *tcsB* | Putative sensor histidine kinase/response regulator with homology to S. cerevisiae Sln1p |
| Afu2g01260 | *srbA* | Sterol regulatory element binding protein (SREBP) |
| Afu2g07770 | *rasB* | Ras family GTPase protein |
| Afu2g12200 | *pkaC* | cAMP-dependent protein kinase catalytic subunit |
| Afu2g12640 | *gprD* | Putative G-protein coupled receptor (GPCR)-like protein |
| Afu2g13260 | *medA* | Putative regulator of adherence, host cell interactions and virulence |
| Afu3g05900 | *ste7* | MAP kinase kinase (MAPKK) |
| Afu3g10000 | *pkaR* | cAMP-dependent protein kinase regulatory subunit |
| Afu3g11080 | *bck1* | Putative mitogen-activated protein kinase kinase kinase (MAPKKK) |
| Afu3g11250 | *ace2* | C2H2 transcription factor with a role in conidiophore development, pigment production, germination and virulence |
| Afu4g13720 | *mpkA* | mitogen-activated protein kinase |
| Afu5g06420 | *steC/ste11* | Ortholog(s) have MAP kinase kinase kinase activity, MAP kinase kinase kinase kinase activity, SAM domain binding activity |
| Afu5g08420 | *sho1* | Putative transmembrane osmosensor with homology to S. cerevisiae Sho1p |
| Afu5g09100 | *mpkC* | Putative mitogen activated protein kinase (MAPK) |
| Afu5g09360 | *calA* | calcineurin a catalytic subunit |
| Afu5g11230 | *rasA* | Ras family GTPase protein |
| Afu5g12210 | *sfaD* | G protein-coupled receptor beta subunit |

| | | |
|---|---|---|
| Afu6g08520 | *acyA* | Adenylate cyclase of the cAMP-dependent signaling pathway, involved in regulation of proliferation and conidiophore development |
| Afu6g10240 | *fos-1* | Putative histidine kinase, two-component signal transduction protein |
| Afu6g12820 | *mpkB* | Putative mitogen-activated protein kinase (MAPK) |
| Afu7g04800 | *gprC* | Rhodopsin-like G-protein coupled receptor |

*null = no gene name assigned.

**Table S3.** Predicted effect of resulting variants after filtering VCF files with SnpSift.

| **Impact and predicted effect** | **Isolates** | | | |
|---|---|---|---|---|
| | **B5233** | **P1MS** | **P1MR** | **P2CS** |
| **High** | 883 | 759 | 877 | 892 |
| Frameshift variant | 346 | 317 | 333 | 333 |
| Splice acceptor | 34 | 24 | 37 | 33 |
| Splice donor | 37 | 29 | 34 | 29 |
| Start lost | 16 | 19 | 19 | 23 |
| Stop gained | 270 | 224 | 269 | 282 |
| Stop lost | 180 | 146 | 185 | 192 |
| **Moderate** | 12,417 | 10,117 | 11,730 | 12,100 |
| Inframe deletion | 173 | 149 | 143 | 168 |
| Inframe insertion | 177 | 164 | 181 | 185 |
| Missense variant | 12,067 | 9,804 | 11,406 | 11,747 |

**TABLE S4.** Examples of variations in virulence related genes.

| Virulence related gene | Isolate | Chromosome number | Position | REF | ALT | Annotation | Annotation_I | Gene_name | Feature_typ | Transcript_E | HGVS.c | HGVS.p | cDNA.pos/c | CDS.pos/CD | AA.pos/AA.length |
|---|---|---|---|---|---|---|---|---|---|---|---|---|---|---|---|
| | B5233 | Chr1_A_fumigatus_Af29 | 1148768 | A | G | missense_variant | MODERATE | Afu1g03992 | transcript | Coding | c.3698T>C | p.Leu1233Pr | 4385/4717 | 3698/3854 | 1233/1283 |
| | | Chr1_A_fumigatus_Af29 | 1149008 | A | G | missense_variant | MODERATE | Afu1g03992 | transcript | Coding | c.3458T>C | p.Met1153T | 4145/4717 | 3458/3854 | 1153/1283 |
| | | Chr1_A_fumigatus_Af29 | 1149189 | G | A | missense_variant | MODERATE | Afu1g03992 | transcript | Coding | c.3277C>T | p.Pro1093Se | 3964/4717 | 3277/3854 | 1093/1283 |
| | P1MR | Chr1_A_fumigatus_Af29 | 1148768 | A | G | missense_variant | MODERATE | Afu1g03992 | transcript | Coding | c.3698T>C | p.Leu1233Pr | 4385/4717 | 3698/3854 | 1233/1283 |
| | | Chr1_A_fumigatus_Af29 | 1149008 | A | G | missense_variant | MODERATE | Afu1g03992 | transcript | Coding | c.3458T>C | p.Met1153T | 4145/4717 | 3458/3854 | 1153/1283 |
| | P1MS | Chr1_A_fumigatus_Af29 | 1148768 | A | G | missense_variant | MODERATE | Afu1g03992 | transcript | Coding | c.3698T>C | p.Leu1233Pr | 4385/4717 | 3698/3854 | 1233/1283 |
| | | Chr1_A_fumigatus_Af29 | 1149008 | A | G | missense_variant | MODERATE | Afu1g03992 | transcript | Coding | c.3458T>C | p.Met1153T | 4145/4717 | 3458/3854 | 1153/1283 |
| | | Chr1_A_fumigatus_Af29 | 1149638 | G | A | missense_variant | MODERATE | Afu1g03992 | transcript | Coding | c.2828C>T | p.Ala943Val | 3515/4717 | 2828/3854 | 943/1283 |
| | | Chr1_A_fumigatus_Af29 | 1150394 | G | A | missense_variant | MODERATE | Afu1g03992 | transcript | Coding | c.2188C>T | p.Leu730Phe | 2875/4717 | 2188/3854 | 730/1283 |
| *thtA* | | Chr1_A_fumigatus_Af29 | 1151572 | C | A | missense_variant | MODERATE | Afu1g03992 | transcript | Coding | c.1010G>T | p.Ser337Ile | 1697/4717 | 1010/3854 | 337/1283 |
| | P2CS | Chr1_A_fumigatus_Af29 | 1148768 | A | G | missense_variant | MODERATE | Afu1g03992 | transcript | Coding | c.3698T>C | p.Leu1233Pr | 4385/4717 | 3698/3854 | 1233/1283 |
| | | Chr1_A_fumigatus_Af29 | 1149008 | A | G | missense_variant | MODERATE | Afu1g03992 | transcript | Coding | c.3458T>C | p.Met1153T | 4145/4717 | 3458/3854 | 1153/1283 |
| | | Chr1_A_fumigatus_Af29 | 1153362 | C | T | 5_prime_UTR_variant | MODIFIER | Afu1g03992 | transcript | Coding | c.-598G>A | | | | |
| | | Chr1_A_fumigatus_Af29 | 1149189 | G | A | missense_variant | MODERATE | Afu1g03992 | transcript | Coding | c.3277C>T | p.Pro1093Se | 3964/4717 | 3277/3854 | 1093/1283 |
| | 08-19-02-30 [ | Chr1_A_fumigatus_Af29 | 1148768 | A | G | missense_variant | MODERATE | Afu1g03992 | transcript | Coding | c.3698T>C | p.Leu1233Pr | 4385/4717 | 3698/3854 | 1153/1283 |
| | | Chr1_A_fumigatus_Af29 | 1149008 | A | G | missense_variant | MODERATE | Afu1g03992 | transcript | Coding | c.3458T>C | p.Met1153T | 4145/4717 | 3458/3854 | 710/1283 |
| | | Chr1_A_fumigatus_Af29 | 1150454 | G | A | stop_gained | HIGH | Afu1g03992 | transcript | Coding | c.2128C>T | p.Arg710* | 2815/4717 | 2128/3854 | 91/1283 |
| | | Chr1_A_fumigatus_Af29 | 1152425 | G | T | missense_variant | MODERATE | Afu1g03992 | transcript | Coding | c.272C>A | p.Ser91Tyr | 959/4717 | 272/3854 | |
| | 08-19-02-46 [ | Chr1_A_fumigatus_Af29 | 1148768 | A | G | missense_variant | MODERATE | Afu1g03992 | transcript | Coding | c.3698T>C | p.Leu1233Pr | 4385/4717 | 3698/3854 | 1153/1283 |
| | | Chr1_A_fumigatus_Af29 | 1149008 | A | G | missense_variant | MODERATE | Afu1g03992 | transcript | Coding | c.3458T>C | p.Met1153T | 4145/4717 | 3458/3854 | |
| | B5233 | Chr3_A_fumigatus_Af29 | 1592149 | A | G | synonymous_variant | LOW | Afu3g06450 | transcript | Coding | c.414A>G | p.Glu138Glu | 743/3170 | 414/2841 | |
| | P1MR | Chr3_A_fumigatus_Af29 | 1597066 | G | A | missense_variant | MODERATE | Afu3g06470 | transcript | Coding | c.442G>A | p.Gly148Ser | 892/2759 | 442/1824 | |
| | P1MS | - | | | | | | | | | | | | | |
| | P2CS | Chr3_A_fumigatus_Af29 | 1591436 | C | T | 5_prime_UTR_variant | MODIFIER | Afu3g06450 | transcript | Coding | c.-232C>T | | | | 775/946 |
| *pmt1* | | Chr3_A_fumigatus_Af29 | 1594250 | C | T | synonymous_variant | LOW | Afu3g06450 | transcript | Coding | c.2325C>T | p.Leu775Leu | 2654/3170 | 2325/2841 | |
| | 08-19-02-30 [ | Chr3_A_fumigatus_Af29 | 1591436 | C | T | 5_prime_UTR_variant | MODIFIER | Afu3g06450 | transcript | Coding | c.-232C>T | | | | 775/946 |
| | | Chr3_A_fumigatus_Af29 | 1594250 | C | T | synonymous_variant | LOW | Afu3g06450 | transcript | Coding | c.2325C>T | p.Leu775Leu | 2654/3170 | 2325/2841 | |
| | 08-19-02-46 [ | Chr3_A_fumigatus_Af29 | 1591919 | C | T | synonymous_variant | LOW | Afu3g06450 | transcript | Coding | c.252C>T | p.Phe84Phe | 581/3170 | 252/2841 | 138/946 |
| | | Chr3_A_fumigatus_Af29 | 1592149 | A | G | synonymous_variant | LOW | Afu3g06450 | transcript | Coding | c.414A>G | p.Glu138Glu | 743/3170 | 414/2841 | |
| | B5233 | Chr1_A_fumigatus_Af29 | 4718702 | T | C | upstream_gene_variant | MODIFIER | Afu1g17250 | transcript | Coding | c.-2771A>G | | | | |
| | | Chr1_A_fumigatus_Af29 | 4719074 | A | G | upstream_gene_variant | MODIFIER | Afu1g17250 | transcript | Coding | c.-3143T>C | | | | |
| | | Chr1_A_fumigatus_Af29 | 4719228 | A | G | upstream_gene_variant | MODIFIER | Afu1g17250 | transcript | Coding | c.-3297T>C | | | | |
| | | Chr1_A_fumigatus_Af29 | 4719412 | TGG | TGGG | upstream_gene_variant | MODIFIER | Afu1g17250 | transcript | Coding | c.-3484_-3483insC | | | | |
| | | Chr1_A_fumigatus_Af29 | 4719456 | C | T | upstream_gene_variant | MODIFIER | Afu1g17250 | transcript | Coding | c.-3525G>A | | | | |
| *rodB* | | Chr1_A_fumigatus_Af29 | 4719642 | C | T | upstream_gene_variant | MODIFIER | Afu1g17250 | transcript | Coding | c.-3711G>A | | | | |
| | | Chr1_A_fumigatus_Af29 | 4719675 | G | C | upstream_gene_variant | MODIFIER | Afu1g17250 | transcript | Coding | c.-3744C>G | | | | |
| | P1MR | - | | | | | | | | | | | | | |
| | P1MS | - | | | | | | | | | | | | | |
| | P2CS | - | | | | | | | | | | | | | |
| | 08-19-02-30 [ | Chr1_A_fumigatus_Af29 | 4719229 | ACCCC | ACCCC | upstream_gene_variant | MODIFIER | Afu1g17250 | transcript | Coding | c.-3308_-3307insG | | | | |
| | 08-19-02-46 [ | Chr1_A_fumigatus_Af29 | 4715857 | A | G | synonymous_variant | LOW | Afu1g17250 | transcript | Coding | c.75T>C | p.Pro25Pro | 588/1221 | 75/552 | |
| | B5233 | Chr3_A_fumigatus_Af29 | 562875 | T | G | 5_prime_UTR_variant | MODIFIER | Afu3g02270 | transcript | Coding | c.-701T>G | | | | |
| | | Chr3_A_fumigatus_Af29 | 562995 | G | A | 5_prime_UTR_variant | MODIFIER | Afu3g02270 | transcript | Coding | c.-581G>A | | | | |
| | | Chr3_A_fumigatus_Af29 | 563334 | T | C | 5_prime_UTR_variant | MODIFIER | Afu3g02270 | transcript | Coding | c.-242T>C | | | | |
| | | Chr3_A_fumigatus_Af29 | 563361 | C | T | 5_prime_UTR_variant | MODIFIER | Afu3g02270 | transcript | Coding | c.-215C>T | | | | 58/728 |
| | | Chr3_A_fumigatus_Af29 | 563749 | C | T | synonymous_variant | LOW | Afu3g02270 | transcript | Coding | c.174C>T | p.Asp58Asp | 1325/3514 | 174/2187 | 148/728 |
| | | Chr3_A_fumigatus_Af29 | 564085 | T | C | synonymous_variant | LOW | Afu3g02270 | transcript | Coding | c.444T>C | p.Gly148Gly | 1595/3514 | 444/2187 | |
| | | Chr3_A_fumigatus_Af29 | 564166 | TT | AC | upstream_gene_variant | MODIFIER | Afu3g02280 | transcript | Coding | c.-2978_-2977delTTinsAC | | | | 207/728 |
| | | Chr3_A_fumigatus_Af29 | 564311 | C | G | synonymous_variant | LOW | Afu3g02270 | transcript | Coding | c.621C>G | p.Ala207Ala | 1772/3514 | 621/2187 | 513/728 |
| | | Chr3_A_fumigatus_Af29 | 565429 | G | A | synonymous_variant | LOW | Afu3g02270 | transcript | Coding | c.1539G>A | p.Val513Val | 2690/3514 | 1539/2187 | |
| | P1MR | Chr3_A_fumigatus_Af29 | 562633 | T | C | 5_prime_UTR_variant | MODIFIER | Afu3g02270 | transcript | Coding | c.-943T>C | | | | |
| | | Chr3_A_fumigatus_Af29 | 562875 | T | G | 5_prime_UTR_variant | MODIFIER | Afu3g02270 | transcript | Coding | c.-701T>G | | | | |
| | | Chr3_A_fumigatus_Af29 | 562921 | G | A | 5_prime_UTR_variant | MODIFIER | Afu3g02270 | transcript | Coding | c.-655G>A | | | | |
| | | Chr3_A_fumigatus_Af29 | 562995 | G | A | 5_prime_UTR_variant | MODIFIER | Afu3g02270 | transcript | Coding | c.-581G>A | | | | |
| | | Chr3_A_fumigatus_Af29 | 563334 | T | C | 5_prime_UTR_variant | MODIFIER | Afu3g02270 | transcript | Coding | c.-242T>C | | | | |
| *cat1* | | Chr3_A_fumigatus_Af29 | 563361 | C | T | 5_prime_UTR_variant | MODIFIER | Afu3g02270 | transcript | Coding | c.-215C>T | | | | 97/728 |

| | | | | | | | | | | | | | | | |
|---|---|---|---|---|---|---|---|---|---|---|---|---|---|---|---|
| *cat1* | | | | | | | | | | | | | | | |
| | | Chr3_A_fumigatus_Af29: | 563932 | C | T | synonymous_variant | LOW | Afu3g02270 | transcript | Coding | c.291C>T | p.Pro97Pro | 1442/3514 | 291/2187 | 148/728 |
| | | Chr3_A_fumigatus_Af29: | 564085 | T | C | synonymous_variant | LOW | Afu3g02270 | transcript | Coding | c.444T>C | p.Gly148Gly | 1595/3514 | 444/2187 | 207/728 |
| | | Chr3_A_fumigatus_Af29: | 564311 | C | G | synonymous_variant | LOW | Afu3g02270 | transcript | Coding | c.621C>G | p.Ala207Ala | 1772/3514 | 621/2187 | 439/728 |
| | | Chr3_A_fumigatus_Af29: | 565207 | C | A | synonymous_variant | LOW | Afu3g02270 | transcript | Coding | c.1317C>A | p.Ser439Ser | 2468/3514 | 1317/2187 | |
| | P1MS | - | | | | | | | | | | | | | |
| | P2CS | - | | | | | | | | | | | | | |
| | 08-19-02-30 D | - | | | | | | | | | | | | | |
| | 08-19-02-46 D | Chr3_A_fumigatus_Af29: | 562875 | T | G | 5_prime_UTR_variant | MODIFIER | Afu3g02270 | transcript | Coding | c.-701T>G | | | | |
| | | Chr3_A_fumigatus_Af29: | 562995 | G | A | 5_prime_UTR_variant | MODIFIER | Afu3g02270 | transcript | Coding | c.-581G>A | | | | |
| | | Chr3_A_fumigatus_Af29: | 563334 | T | C | 5_prime_UTR_variant | MODIFIER | Afu3g02270 | transcript | Coding | c.-242T>C | | | | |
| | | Chr3_A_fumigatus_Af29: | 563361 | C | T | 5_prime_UTR_variant | MODIFIER | Afu3g02270 | transcript | Coding | c.-215C>T | | | | 58/728 |
| | | Chr3_A_fumigatus_Af29: | 563749 | C | T | synonymous_variant | LOW | Afu3g02270 | transcript | Coding | c.174C>T | p.Asp58Asp | 1325/3514 | 174/2187 | 148/728 |
| | | Chr3_A_fumigatus_Af29: | 564085 | T | C | synonymous_variant | LOW | Afu3g02270 | transcript | Coding | c.444T>C | p.Gly148Gly | 1595/3514 | 444/2187 | 207/728 |
| | | Chr3_A_fumigatus_Af29: | 564311 | C | G | synonymous_variant | LOW | Afu3g02270 | transcript | Coding | c.621C>G | p.Ala207Ala | 1772/3514 | 621/2187 | 513/728 |
| | | Chr3_A_fumigatus_Af29: | 565429 | G | A | synonymous_variant | LOW | Afu3g02270 | transcript | Coding | c.1539G>A | p.Val513Val | 2690/3514 | 1539/2187 | |
| | B5233 | Chr6_A_fumigatus_Af29: | 855633 | C | T | 5_prime_UTR_variant | MODIFIER | Afu6g03890 | transcript | Coding | c.-1243C>T | | | | |
| | | Chr6_A_fumigatus_Af29: | 856139 | G | A | 5_prime_UTR_variant | MODIFIER | Afu6g03890 | transcript | Coding | c.-737G>A | | | | |
| | | Chr6_A_fumigatus_Af29: | 856407 | G | A | 5_prime_UTR_variant | MODIFIER | Afu6g03890 | transcript | Coding | c.-469G>A | | | | |
| | | Chr6_A_fumigatus_Af29: | 856418 | C | T | 5_prime_UTR_variant | MODIFIER | Afu6g03890 | transcript | Coding | c.-458C>T | | | | |
| | | Chr6_A_fumigatus_Af29: | 856713 | A | T | 5_prime_UTR_variant | MODIFIER | Afu6g03890 | transcript | Coding | c.-163A>T | | | | |
| | | Chr6_A_fumigatus_Af29: | 859250 | A | T | 3_prime_UTR_variant | MODIFIER | Afu6g03890 | transcript | Coding | c.*16A>T | | | | |
| | P1MR | Chr6_A_fumigatus_Af29: | 856139 | G | A | 5_prime_UTR_variant | MODIFIER | Afu6g03890 | transcript | Coding | c.-737G>A | | | | |
| | | Chr6_A_fumigatus_Af29: | 856407 | G | A | 5_prime_UTR_variant | MODIFIER | Afu6g03890 | transcript | Coding | c.-469G>A | | | | |
| | | Chr6_A_fumigatus_Af29: | 856685 | C | T | 5_prime_UTR_premature_st | LOW | Afu6g03890 | transcript | Coding | c.-191C>T | | | | |
| | | Chr6_A_fumigatus_Af29: | 856713 | A | T | 5_prime_UTR_variant | MODIFIER | Afu6g03890 | transcript | Coding | c.-163A>T | | | | 462/750 |
| | | Chr6_A_fumigatus_Af29: | 858366 | G | A | missense_variant | MODERATE | Afu6g03890 | transcript | Coding | c.1385G>A | p.Ser462Asn | 3291/4243 | 1385/2253 | |
| | P1MS | Chr6_A_fumigatus_Af29: | 857963 | G | A | missense_variant | MODERATE | Afu6g03890 | transcript | Coding | c.982G>A | p.Asp328Asn | 2888/4243 | 982/2253 | 462/750 |
| | | Chr6_A_fumigatus_Af29: | 858366 | G | A | missense_variant | MODERATE | Afu6g03890 | transcript | Coding | c.1385G>A | p.Ser462Asn | 3291/4243 | 1385/2253 | 690/1051 |
| *catA* | P2CS | Chr6_A_fumigatus_Af29: | 855635 | C | G | 5_prime_UTR_variant | MODIFIER | Afu6g03890 | transcript | Coding | c.-1241C>G | | | | |
| | | Chr6_A_fumigatus_Af29: | 856139 | G | A | 5_prime_UTR_variant | MODIFIER | Afu6g03890 | transcript | Coding | c.-737G>A | | | | |
| | | Chr6_A_fumigatus_Af29: | 856407 | G | A | 5_prime_UTR_variant | MODIFIER | Afu6g03890 | transcript | Coding | c.-469G>A | | | | |
| | | Chr6_A_fumigatus_Af29: | 856713 | A | T | 5_prime_UTR_variant | MODIFIER | Afu6g03890 | transcript | Coding | c.-163A>T | | | | 381/750 |
| | | Chr6_A_fumigatus_Af29: | 858124 | C | T | synonymous_variant | LOW | Afu6g03890 | transcript | Coding | c.1143C>T | p.Phe381Phe | 3049/4243 | 1143/2253 | |
| | 08-19-02-30 D | Chr6_A_fumigatus_Af29: | 856139 | G | A | 5_prime_UTR_variant | MODIFIER | Afu6g03890 | transcript | Coding | c.-737G>A | | | | |
| | | Chr6_A_fumigatus_Af29: | 856407 | G | A | 5_prime_UTR_variant | MODIFIER | Afu6g03890 | transcript | Coding | c.-469G>A | | | | |
| | | Chr6_A_fumigatus_Af29: | 856713 | A | T | 5_prime_UTR_variant | MODIFIER | Afu6g03890 | transcript | Coding | c.-163A>T | | | | 381/750 |
| | | Chr6_A_fumigatus_Af29: | 858124 | C | T | synonymous_variant | LOW | Afu6g03890 | transcript | Coding | c.1143C>T | p.Phe381Phe | 3049/4243 | 1143/2253 | |
| | 08-19-02-46 D | Chr6_A_fumigatus_Af29: | 855635 | C | G | 5_prime_UTR_variant | MODIFIER | Afu6g03890 | transcript | Coding | c.-1241C>G | | | | |
| | | Chr6_A_fumigatus_Af29: | 856139 | G | A | 5_prime_UTR_variant | MODIFIER | Afu6g03890 | transcript | Coding | c.-737G>A | | | | |
| | | Chr6_A_fumigatus_Af29: | 856407 | G | A | 5_prime_UTR_variant | MODIFIER | Afu6g03890 | transcript | Coding | c.-469G>A | | | | |
| | | Chr6_A_fumigatus_Af29: | 856713 | A | T | 5_prime_UTR_variant | MODIFIER | Afu6g03890 | transcript | Coding | c.-163A>T | | | | 381/750 |
| | | Chr6_A_fumigatus_Af29: | 858124 | C | T | synonymous_variant | LOW | Afu6g03890 | transcript | Coding | c.1143C>T | p.Phe381Phe | 3049/4243 | 1143/2253 | |
| | B5233 | Chr1_A_fumigatus_Af29: | 2169850 | C | T | synonymous_variant | LOW | Afu1g07690 | transcript | Coding | c.990G>A | p.Gln330Gln | 1169/2721 | 990/2250 | |
| | P1MR | Chr1_A_fumigatus_Af29: | 2169850 | C | T | synonymous_variant | LOW | Afu1g07690 | transcript | Coding | c.990G>A | p.Gln330Gln | 1169/2721 | 990/2250 | |
| *afpmt2* | P1MS | - | | | | | | | | | | | | | |
| | P2CS | - | | | | | | | | | | | | | 480/763 |
| | 08-19-02-30 D | - | | | | | | | | | | | | | |
| | 08-19-02-46 D | Chr1_A_fumigatus_Af29: | 2169850 | C | T | synonymous_variant | LOW | Afu1g07690 | transcript | Coding | c.990G>A | p.Gln330Gln | 1169/2721 | 990/2250 | |
| | B5233 | - | | | | | | | | | | | | | |
| | P1MR | - | | | | | | | | | | | | | |
| | P1MS | Chr1_A_fumigatus_Af29: | 3918209 | G | A | stop_gained | HIGH | Afu1g14660 | transcript | Coding | c.189G>A | p.Trp63* | 189/1122 | 189/1122 | 134/373 |
| *laeA* | | Chr1_A_fumigatus_Af29: | 3918565 | C | T | missense_variant | MODERATE | Afu1g14660 | transcript | Coding | c.400C>T | p.Pro134Ser | 400/1122 | 400/1122 | |
| | P2CS | - | | | | | | | | | | | | | |
| | 08-19-02-30 D | - | | | | | | | | | | | | | |
| | 08-19-02-46 D | - | | | | | | | | | | | | | |
| | B5233 | Chr6_A_fumigatus_Af29: | 2347394 | GCAAC | GCAAC | disruptive_inframe_deletion | MODERATE | Afu6g09630 | transcript | Coding | c.425_427de | p.Thr142del | 898/1943 | 425/1470 | 142/489 |
| | | Chr6_A_fumigatus_Af29: | 2348162 | C | G | missense_variant | MODERATE | Afu6g09630 | transcript | Coding | c.1177C>G | p.Leu393Val | 1650/1943 | 1177/1470 | 393/489 |
| | P1MR | Chr6_A_fumigatus_Af29: | 2346580 | T | C | 5_prime_UTR_variant | MODIFIER | Afu6g09630 | transcript | Coding | c.-406T>C | | | | |

| | | | | | | | | | | | | | | | |
|---|---|---|---|---|---|---|---|---|---|---|---|---|---|---|---|
| | | Chr6_A_fumigatus_Af29: | 2346745 | T | C | 5_prime_UTR_variant | MODIFIER | Afu6g09630 | transcript | Coding | c.-241T>C | | | | |
| | | Chr6_A_fumigatus_Af29: | 2346807 | G | C | 5_prime_UTR_variant | MODIFIER | Afu6g09630 | transcript | Coding | c.-179G>C | | | | |
| | | Chr6_A_fumigatus_Af29: | 2346831 | C | T | 5_prime_UTR_variant | MODIFIER | Afu6g09630 | transcript | Coding | c.-155C>T | | | | |
| | | Chr6_A_fumigatus_Af29: | 2346890 | A | C | 5_prime_UTR_variant | MODIFIER | Afu6g09630 | transcript | Coding | c.-96A>C | | | | |
| | | Chr6_A_fumigatus_Af29: | 2346922 | A | C | 5_prime_UTR_variant | MODIFIER | Afu6g09630 | transcript | Coding | c.-64A>C | | | | 27/489 |
| | | Chr6_A_fumigatus_Af29: | 2347064 | A | G | missense_variant | MODERATE | Afu6g09630 | transcript | Coding | c.79A>G | p.Thr27Ala | 552/1943 | 79/1470 | 34/489 |
| | | Chr6_A_fumigatus_Af29: | 2347073 | ACTGC | ACTGC | disruptive_inframe_insertion | MODERATE | Afu6g09630 | transcript | Coding | c.99_101dup | p.Ala34dup | 575/1943 | 102/1470 | 108/489 |
| | | Chr6_A_fumigatus_Af29: | 2347309 | A | C | synonymous_variant | LOW | Afu6g09630 | transcript | Coding | c.324A>C | p.Ile108Ile | 797/1943 | 324/1470 | 130/489 |
| | | Chr6_A_fumigatus_Af29: | 2347373 | A | G | missense_variant | MODERATE | Afu6g09630 | transcript | Coding | c.388A>G | p.Thr130Ala | 861/1943 | 388/1470 | 136/489 |
| | | Chr6_A_fumigatus_Af29: | 2347388 | GC | GCAAC | inframe_insertion | MODERATE | Afu6g09630 | transcript | Coding | c.405_406in | p.Ala135_Al | 879/1943 | 406/1470 | 136/489 |
| | | Chr6_A_fumigatus_Af29: | 2347391 | GCAG | ACAA | missense_variant | MODERATE | Afu6g09630 | transcript | Coding | c.406_409de | p.AlaAla136 | 879/1943 | 406/1470 | 155/489 |
| | | Chr6_A_fumigatus_Af29: | 2347449 | T | G | missense_variant | MODERATE | Afu6g09630 | transcript | Coding | c.464T>G | p.Val155Gly | 937/1943 | 464/1470 | 204/489 |
| | | Chr6_A_fumigatus_Af29: | 2347597 | C | T | synonymous_variant | LOW | Afu6g09630 | transcript | Coding | c.612C>T | p.Ser204Ser | 1085/1943 | 612/1470 | 240/489 |
| | | Chr6_A_fumigatus_Af29: | 2347703 | T | G | missense_variant | MODERATE | Afu6g09630 | transcript | Coding | c.718T>G | p.Trp240Gly | 1191/1943 | 718/1470 | 363/489 |
| | | Chr6_A_fumigatus_Af29: | 2348072 | T | C | missense_variant | MODERATE | Afu6g09630 | transcript | Coding | c.1087T>C | p.Phe363Leu | 1560/1943 | 1087/1470 | |
| | P1MS | Chr6_A_fumigatus_Af29: | 2347373 | A | G | missense_variant | MODERATE | Afu6g09630 | transcript | Coding | c.388A>G | p.Thr130Ala | 861/1943 | 388/1470 | 133/489 |
| | | Chr6_A_fumigatus_Af29: | 2347382 | GCAGC | ACAAC | missense_variant&disruptive | MODERATE | Afu6g09630 | transcript | Coding | c.397_409de | p.Ala133_Al | 870/1943 | 397/1470 | 155/489 |
| | | Chr6_A_fumigatus_Af29: | 2347449 | T | G | missense_variant | MODERATE | Afu6g09630 | transcript | Coding | c.464T>G | p.Val155Gly | 937/1943 | 464/1470 | 363/489 |
| | | Chr6_A_fumigatus_Af29: | 2348072 | T | C | missense_variant | MODERATE | Afu6g09630 | transcript | Coding | c.1087T>C | p.Phe363Leu | 1560/1943 | 1087/1470 | |
| | P2CS | Chr6_A_fumigatus_Af29: | 2346580 | T | C | 5_prime_UTR_variant | MODIFIER | Afu6g09630 | transcript | Coding | c.-406T>C | | | | |
| | | Chr6_A_fumigatus_Af29: | 2346745 | T | C | 5_prime_UTR_variant | MODIFIER | Afu6g09630 | transcript | Coding | c.-241T>C | | | | |
| *gliZ* | | Chr6_A_fumigatus_Af29: | 2346807 | G | C | 5_prime_UTR_variant | MODIFIER | Afu6g09630 | transcript | Coding | c.-179G>C | | | | |
| | | Chr6_A_fumigatus_Af29: | 2346890 | A | C | 5_prime_UTR_variant | MODIFIER | Afu6g09630 | transcript | Coding | c.-96A>C | | | | |
| | | Chr6_A_fumigatus_Af29: | 2346922 | A | C | 5_prime_UTR_variant | MODIFIER | Afu6g09630 | transcript | Coding | c.-64A>C | | | | |
| | | Chr6_A_fumigatus_Af29: | 2347073 | ACTGC | ACTGC | disruptive_inframe_insertion | MODERATE | Afu6g09630 | transcript | Coding | c.99_101dup | p.Ala34dup | 575/1943 | 102/1470 | 34/489 |
| | | Chr6_A_fumigatus_Af29: | 2347309 | A | C | synonymous_variant | LOW | Afu6g09630 | transcript | Coding | c.324A>C | p.Ile108Ile | 797/1943 | 324/1470 | 108/489 |
| | | Chr6_A_fumigatus_Af29: | 2347373 | A | G | missense_variant | MODERATE | Afu6g09630 | transcript | Coding | c.388A>G | p.Thr130Ala | 861/1943 | 388/1470 | 130/489 |
| | | Chr6_A_fumigatus_Af29: | 2347394 | GC | GCAGC | inframe_insertion | MODERATE | Afu6g09630 | transcript | Coding | c.411_412in | p.Ala137_Th | 885/1943 | 412/1470 | 138/489 |
| | | Chr6_A_fumigatus_Af29: | 2347397 | A | G | missense_variant | MODERATE | Afu6g09630 | transcript | Coding | c.412A>G | p.Thr138Ala | 885/1943 | 412/1470 | 138/489 |
| | | Chr6_A_fumigatus_Af29: | 2347449 | T | G | missense_variant | MODERATE | Afu6g09630 | transcript | Coding | c.464T>G | p.Val155Gly | 937/1943 | 464/1470 | 155/489 |
| | | Chr6_A_fumigatus_Af29: | 2347597 | C | T | synonymous_variant | LOW | Afu6g09630 | transcript | Coding | c.612C>T | p.Ser204Ser | 1085/1943 | 612/1470 | 204/489 |
| | | Chr6_A_fumigatus_Af29: | 2348072 | T | C | missense_variant | MODERATE | Afu6g09630 | transcript | Coding | c.1087T>C | p.Phe363Leu | 1560/1943 | 1087/1470 | 363/489 |
| | 08-19-02-30 D | Chr6_A_fumigatus_Af29: | 2346648 | CAA | CAAA | 5_prime_UTR_variant | MODIFIER | Afu6g09630 | transcript | Coding | c.-334dupA | | | | 119/489 |
| | | Chr6_A_fumigatus_Af29: | 2347342 | G | A | synonymous_variant | LOW | Afu6g09630 | transcript | Coding | c.357G>A | p.Pro119Pro | 830/1943 | 357/1470 | 130/489 |
| | | Chr6_A_fumigatus_Af29: | 2347373 | A | G | missense_variant | MODERATE | Afu6g09630 | transcript | Coding | c.388A>G | p.Thr130Ala | 861/1943 | 388/1470 | 133/489 |
| | | Chr6_A_fumigatus_Af29: | 2347382 | GCAGC | ACAAC | missense_variant&disruptive | MODERATE | Afu6g09630 | transcript | Coding | c.397_409de | p.Ala133_Al | 870/1943 | 397/1470 | 155/489 |
| | | Chr6_A_fumigatus_Af29: | 2347449 | T | G | missense_variant | MODERATE | Afu6g09630 | transcript | Coding | c.464T>G | p.Val155Gly | 937/1943 | 464/1470 | 187/489 |
| | | Chr6_A_fumigatus_Af29: | 2347546 | G | A | synonymous_variant | LOW | Afu6g09630 | transcript | Coding | c.561G>A | p.Ser187Ser | 1034/1943 | 561/1470 | 204/489 |
| | | Chr6_A_fumigatus_Af29: | 2347597 | C | T | synonymous_variant | LOW | Afu6g09630 | transcript | Coding | c.612C>T | p.Ser204Ser | 1085/1943 | 612/1470 | 446/489 |
| | | Chr6_A_fumigatus_Af29: | 2348305 | GTCCTC | GTCCT | disruptive_inframe_deletion | MODERATE | Afu6g09630 | transcript | Coding | c.1338_1340 | p.Ser447del | 1811/1943 | 1338/1470 | 479/489 |
| | | Chr6_A_fumigatus_Af29: | 2348420 | A | G | missense_variant | MODERATE | Afu6g09630 | transcript | Coding | c.1435A>G | p.Ile479Val | 1908/1943 | 1435/1470 | |
| | 512 08-19-02- | Chr6_A_fumigatus_Af29: | 2347373 | ACAGC | ACAGC | inframe_insertion | MODERATE | Afu6g09630 | transcript | Coding | c.406_411du | p.Ala136_Al | 885/1943 | 412/1470 | 138/489 |
| | | Chr6_A_fumigatus_Af29: | 2347397 | A | G | missense_variant | MODERATE | Afu6g09630 | transcript | Coding | c.412A>G | p.Thr138Ala | 885/1943 | 412/1470 | 139/489 |
| | | Chr6_A_fumigatus_Af29: | 2347400 | A | G | missense_variant | MODERATE | Afu6g09630 | transcript | Coding | c.415A>G | p.Thr139Ala | 888/1943 | 415/1470 | 393/489 |
| | | Chr6_A_fumigatus_Af29: | 2348162 | C | G | missense_variant | MODERATE | Afu6g09630 | transcript | Coding | c.1177C>G | p.Leu393Val | 1650/1943 | 1177/1470 | 468/489 |
| | | Chr6_A_fumigatus_Af29: | 2348389 | A | C | missense_variant | MODERATE | Afu6g09630 | transcript | Coding | c.1404A>C | p.Glu468Asp | 1877/1943 | 1404/1470 | |
| | B5233 | Chr1_A_fumigatus_Af29: | 3890479 | G | C | missense_variant | MODERATE | Afu1g14560 | transcript | Coding | c.328C>G | p.Gln110Glu | 927/2176 | 328/1510 | 110/502 |
| | | Chr1_A_fumigatus_Af29: | 3890512 | A | G | stop_lost | HIGH | Afu1g14560 | transcript | Coding | c.295T>C | p.Ter99Glne | 894/2176 | 295/1510 | 99/502 |
| | | Chr1_A_fumigatus_Af29: | 3890599 | A | T | missense_variant | MODERATE | Afu1g14560 | transcript | Coding | c.208T>A | p.Leu70Met | 807/2176 | 208/1510 | 70/502 |
| | | Chr1_A_fumigatus_Af29: | 3891444 | TGACT | GGACC | 5_prime_UTR_variant | MODIFIER | Afu1g14560 | transcript | Coding | c.-580_-576delAGTCAinsGGTCC | | | | |
| | | Chr1_A_fumigatus_Af29: | 3891500 | AC | CT | upstream_gene_variant | MODIFIER | Afu1g14540 | transcript | Coding | c.-4968_-4967delGTinsAG | | | | |
| | | Chr1_A_fumigatus_Af29: | 3891570 | T | A | upstream_gene_variant | MODIFIER | Afu1g14560 | transcript | Coding | c.-702A>T | | | | |
| | | Chr1_A_fumigatus_Af29: | 3892362 | A | C | 5_prime_UTR_variant | MODIFIER | Afu1g14570 | transcript | Coding | c.-191A>C | | | | |
| | | Chr1_A_fumigatus_Af29: | 3893965 | G | A | synonymous_variant | LOW | Afu1g14570 | transcript | Coding | c.1413G>A | p.Val471Val | 1963/3200 | 1413/2619 | 471/872 |
| | | Chr1_A_fumigatus_Af29: | 3895228 | GTTTTT | GTTTT | splice_region_variant | LOW | Afu1g14570 | transcript | Coding | c.*28_*33delTTTTTT | | | | |
| | | Chr1_A_fumigatus_Af29: | 3895252 | G | T | upstream_gene_variant | MODIFIER | Afu1g14560 | transcript | Coding | c.-4384C>A | | | | |
| | | Chr1_A_fumigatus_Af29: | 3895365 | T | G | upstream_gene_variant | MODIFIER | Afu1g14560 | transcript | Coding | c.-4497A>C | | | | |
| | | Chr1_A_fumigatus_Af29: | 3895498 | C | T | upstream_gene_variant | MODIFIER | Afu1g14560 | transcript | Coding | c.-4630G>A | | | | |

| | | | | | | | | | | | | | | | |
|---|---|---|---|---|---|---|---|---|---|---|---|---|---|---|---|
| | P1MR | Chr1_A_fumigatus_Af29 | 3890512 | A | G | stop_lost | HIGH | Afu1g14560 | transcript | Coding | c.295T>C | p.Ter99Glne | 894/2176 | 295/1510 | 99/502 |
| | | Chr1_A_fumigatus_Af29 | 3891181 | C | T | 5_prime_UTR_variant | MODIFIER | Afu1g14560 | transcript | Coding | c.-313G>A | | | | |
| | | Chr1_A_fumigatus_Af29 | 3895380 | T | C | upstream_gene_variant | MODIFIER | Afu1g14560 | transcript | Coding | c.-4512A>G | | | | |
| | P1MS | Chr1_A_fumigatus_Af29 | 3890479 | G | C | missense_variant | MODERATE | Afu1g14560 | transcript | Coding | c.328C>G | p.Gln110Glu | 927/2176 | 328/1510 | 110/502 |
| | | Chr1_A_fumigatus_Af29 | 3890512 | A | G | stop_lost | HIGH | Afu1g14560 | transcript | Coding | c.295T>C | p.Ter99Glne | 894/2176 | 295/1510 | 99/502 |
| | | Chr1_A_fumigatus_Af29 | 3890599 | A | T | missense_variant | MODERATE | Afu1g14560 | transcript | Coding | c.208T>A | p.Leu70Met | 807/2176 | 208/1510 | 70/502 |
| *msdS* | P2CS | Chr1_A_fumigatus_Af29 | 3890479 | G | C | missense_variant | MODERATE | Afu1g14560 | transcript | Coding | c.328C>G | p.Gln110Glu | 927/2176 | 328/1510 | 110/502 |
| | | Chr1_A_fumigatus_Af29 | 3890512 | A | G | stop_lost | HIGH | Afu1g14560 | transcript | Coding | c.295T>C | p.Ter99Glne | 894/2176 | 295/1510 | 99/502 |
| | | Chr1_A_fumigatus_Af29 | 3890599 | A | T | missense_variant | MODERATE | Afu1g14560 | transcript | Coding | c.208T>A | p.Leu70Met | 807/2176 | 208/1510 | 70/502 |
| | | Chr1_A_fumigatus_Af29 | 3891444 | TGACT | GGACC | 5_prime_UTR_variant | MODIFIER | Afu1g14560 | transcript | Coding | c.-580_-576delAGTCAinsGGTCC | | | | |
| | | Chr1_A_fumigatus_Af29 | 3891570 | T | A | upstream_gene_variant | MODIFIER | Afu1g14560 | transcript | Coding | c.-702A>T | | | | |
| | | Chr1_A_fumigatus_Af29 | 3895252 | G | T | upstream_gene_variant | MODIFIER | Afu1g14560 | transcript | Coding | c.-4384C>A | | | | |
| | | Chr1_A_fumigatus_Af29 | 3895365 | T | G | upstream_gene_variant | MODIFIER | Afu1g14560 | transcript | Coding | c.-4497A>C | | | | |
| | | Chr1_A_fumigatus_Af29 | 3895498 | C | T | upstream_gene_variant | MODIFIER | Afu1g14560 | transcript | Coding | c.-4630G>A | | | | |
| | 08-19-02-30 D | Chr1_A_fumigatus_Af29 | 3890512 | A | G | stop_lost | HIGH | Afu1g14560 | transcript | Coding | c.295T>C | p.Ter99Glne | 894/2176 | 295/1510 | |
| | | Chr1_A_fumigatus_Af29 | 3891670 | CTATG | CC | upstream_gene_variant | MODIFIER | Afu1g14560 | transcript | Coding | c.-813_-803delAGTGATGCATA | | | | |
| | 08-19-02-46 | Dutch envornmenatal | | | | | | | | | | | | | 348/502 |
| | | Chr1_A_fumigatus_Af29 | 3889764 | G | A | missense_variant | MODERATE | Afu1g14560 | transcript | Coding | c.1043C>T | p.Ala348Val | 1642/2176 | 1043/1510 | 323/502 |
| | | Chr1_A_fumigatus_Af29 | 3889840 | G | A | synonymous_variant | LOW | Afu1g14560 | transcript | Coding | c.967C>T | p.Leu323Leu | 1566/2176 | 967/1510 | 110/502 |
| | | Chr1_A_fumigatus_Af29 | 3890479 | G | C | missense_variant | MODERATE | Afu1g14560 | transcript | Coding | c.328C>G | p.Gln110Glu | 927/2176 | 328/1510 | 99/502 |
| | | Chr1_A_fumigatus_Af29 | 3890512 | A | G | stop_lost | HIGH | Afu1g14560 | transcript | Coding | c.295T>C | p.Ter99Glne | 894/2176 | 295/1510 | 70/502 |
| | | Chr1_A_fumigatus_Af29 | 3890599 | A | T | missense_variant | MODERATE | Afu1g14560 | transcript | Coding | c.208T>A | p.Leu70Met | 807/2176 | 208/1510 | |
| | | Chr1_A_fumigatus_Af29 | 3891570 | T | A | upstream_gene_variant | MODIFIER | Afu1g14560 | transcript | Coding | c.-702A>T | | | | |
| | | Chr1_A_fumigatus_Af29 | 3895252 | G | T | upstream_gene_variant | MODIFIER | Afu1g14560 | transcript | Coding | c.-4384C>A | | | | |
| | | Chr1_A_fumigatus_Af29 | 3895365 | T | G | upstream_gene_variant | MODIFIER | Afu1g14560 | transcript | Coding | c.-4497A>C | | | | |
| | | Chr1_A_fumigatus_Af29 | 3895498 | C | T | upstream_gene_variant | MODIFIER | Afu1g14560 | transcript | Coding | c.-4630G>A | | | | |
| | B5233 | Chr1_A_fumigatus_Af29 | 4688032 | A | C | 5_prime_UTR_variant | MODIFIER | Afu1g17200 | transcript | Coding | c.-706A>C | | | | |
| | | Chr1_A_fumigatus_Af29 | 4688042 | T | C | 5_prime_UTR_variant | MODIFIER | Afu1g17200 | transcript | Coding | c.-696T>C | | | | 193/4763 |
| | | Chr1_A_fumigatus_Af29 | 4689426 | A | G | missense_variant | MODERATE | Afu1g17200 | transcript | Coding | c.577A>G | p.Ile193Val | 1492/15449 | 577/14292 | 398/4763 |
| | | Chr1_A_fumigatus_Af29 | 4690043 | A | T | synonymous_variant | LOW | Afu1g17200 | transcript | Coding | c.1194A>T | p.Leu398Leu | 2109/15449 | 1194/14292 | 470/4763 |
| | | Chr1_A_fumigatus_Af29 | 4690259 | A | G | synonymous_variant | LOW | Afu1g17200 | transcript | Coding | c.1410A>G | p.Thr470Thr | 2325/15449 | 1410/14292 | 523/4763 |
| | | Chr1_A_fumigatus_Af29 | 4690418 | C | G | missense_variant | MODERATE | Afu1g17200 | transcript | Coding | c.1569C>G | p.Ser523Arg | 2484/15449 | 1569/14292 | 771/4763 |
| | | Chr1_A_fumigatus_Af29 | 4691160 | G | A | missense_variant | MODERATE | Afu1g17200 | transcript | Coding | c.2311G>A | p.Ala771Thr | 3226/15449 | 2311/14292 | 1131/4763 |
| | | Chr1_A_fumigatus_Af29 | 4692240 | A | G | missense_variant | MODERATE | Afu1g17200 | transcript | Coding | c.3391A>G | p.Asn1131A | 4306/15449 | 3391/14292 | 1274/4763 |
| | | Chr1_A_fumigatus_Af29 | 4692669 | A | G | missense_variant | MODERATE | Afu1g17200 | transcript | Coding | c.3820A>G | p.Ile1274Val | 4735/15449 | 3820/14292 | 1917/4763 |
| | | Chr1_A_fumigatus_Af29 | 4694600 | A | G | synonymous_variant | LOW | Afu1g17200 | transcript | Coding | c.5751A>G | p.Ala1917Al | 6666/15449 | 5751/14292 | 2161/4763 |
| | | Chr1_A_fumigatus_Af29 | 4695332 | C | T | synonymous_variant | LOW | Afu1g17200 | transcript | Coding | c.6483C>T | p.Phe2161Pl | 7398/15449 | 6483/14292 | 2242/4763 |
| | | Chr1_A_fumigatus_Af29 | 4695575 | G | A | synonymous_variant | LOW | Afu1g17200 | transcript | Coding | c.6726G>A | p.Arg2242Ar | 7641/15449 | 6726/14292 | 2392/4763 |
| | | Chr1_A_fumigatus_Af29 | 4696023 | A | G | missense_variant | MODERATE | Afu1g17200 | transcript | Coding | c.7174A>G | p.Lys2392Gl | 8089/15449 | 7174/14292 | 3200/4763 |
| | | Chr1_A_fumigatus_Af29 | 4698447 | G | A | missense_variant | MODERATE | Afu1g17200 | transcript | Coding | c.9598G>A | p.Gly3200Se | 10513/1544 | 9598/14292 | 3234/4763 |
| | | Chr1_A_fumigatus_Af29 | 4698551 | C | G | synonymous_variant | LOW | Afu1g17200 | transcript | Coding | c.9702C>G | p.Leu3234Le | 10617/1544 | 9702/14292 | 3243/4763 |
| | | Chr1_A_fumigatus_Af29 | 4698576 | T | C | missense_variant | MODERATE | Afu1g17200 | transcript | Coding | c.9727T>C | p.Phe3243Le | 10642/1544 | 9727/14292 | 3776/4763 |
| | | Chr1_A_fumigatus_Af29 | 4700175 | A | G | missense_variant | MODERATE | Afu1g17200 | transcript | Coding | c.11326A>G | p.Thr3776Al | 12241/1544 | 11326/1429 | 3940/4763 |
| | | Chr1_A_fumigatus_Af29 | 4700669 | T | A | synonymous_variant | LOW | Afu1g17200 | transcript | Coding | c.11820T>A | p.Ala3940Al | 12735/1544 | 11820/1429 | 3979/4763 |
| | | Chr1_A_fumigatus_Af29 | 4700784 | T | G | missense_variant | MODERATE | Afu1g17200 | transcript | Coding | c.11935T>G | p.Ser3979Al | 12850/1544 | 11935/1429 | 4475/4763 |
| | | Chr1_A_fumigatus_Af29 | 4702274 | T | C | synonymous_variant | LOW | Afu1g17200 | transcript | Coding | c.13425T>C | p.Ser4475Se | 14340/1544 | 13425/1429 | 4741/4763 |
| | | Chr1_A_fumigatus_Af29 | 4703071 | G | A | missense_variant | MODERATE | Afu1g17200 | transcript | Coding | c.14222G>A | p.Gly4741Gl | 15137/1544 | 14222/14292 | |
| | | Chr1_A_fumigatus_Af29 | 4703487 | G | A | downstream_gene_variant | MODIFIER | Afu1g17200 | transcript | Coding | c.*346G>A | | | | |
| | | Chr1_A_fumigatus_Af29 | 4703627 | C | A | downstream_gene_variant | MODIFIER | Afu1g17200 | transcript | Coding | c.*486C>A | | | | |
| | | Chr1_A_fumigatus_Af29 | 4703703 | CAA | CTAG | downstream_gene_variant | MODIFIER | Afu1g17200 | transcript | Coding | c.*563_*564delAAinsTAG | | | | |
| | | Chr1_A_fumigatus_Af29 | 4703980 | GCTGG | GCTGG | downstream_gene_variant | MODIFIER | Afu1g17200 | transcript | Coding | c.*852_*862delTGGAAAAAGTC | | | | |
| | | Chr1_A_fumigatus_Af29 | 4704487 | C | G | downstream_gene_variant | MODIFIER | Afu1g17200 | transcript | Coding | c.*1346C>G | | | | |
| | P1MR | Chr1_A_fumigatus_Af29 | 4687837 | T | G | 5_prime_UTR_variant | MODIFIER | Afu1g17200 | transcript | Coding | c.-901T>G | | | | |
| | | Chr1_A_fumigatus_Af29 | 4688032 | A | C | 5_prime_UTR_variant | MODIFIER | Afu1g17200 | transcript | Coding | c.-706A>C | | | | |
| | | Chr1_A_fumigatus_Af29 | 4688042 | T | C | 5_prime_UTR_variant | MODIFIER | Afu1g17200 | transcript | Coding | c.-696T>C | | | | 193/4763 |
| | | Chr1_A_fumigatus_Af29 | 4689426 | A | G | missense_variant | MODERATE | Afu1g17200 | transcript | Coding | c.577A>G | p.Ile193Val | 1492/15449 | 577/14292 | 470/4763 |
| | | Chr1_A_fumigatus_Af29 | 4690259 | A | G | synonymous_variant | LOW | Afu1g17200 | transcript | Coding | c.1410A>G | p.Thr470Thr | 2325/15449 | 1410/14292 | 523/4763 |
| | | Chr1_A_fumigatus_Af29 | 4690418 | C | G | missense_variant | MODERATE | Afu1g17200 | transcript | Coding | c.1569C>G | p.Ser523Arg | 2484/15449 | 1569/14292 | 771/4763 |

| | | | | | | | | | | | | | | | |
|---|---|---|---|---|---|---|---|---|---|---|---|---|---|---|---|
| | | Chr1_A_fumigatus_Af29 | 4691160 | G | A | missense_variant | MODERATE | Afu1g17200 | transcript | Coding | c.2311G>A | p.Ala771Thr | 3226/15449 | 2311/14292 | 1131/4763 |
| | | Chr1_A_fumigatus_Af29 | 4692240 | A | G | missense_variant | MODERATE | Afu1g17200 | transcript | Coding | c.3391A>G | p.Asn1131A | 4306/15449 | 3391/14292 | 1134/4763 |
| | | Chr1_A_fumigatus_Af29 | 4692250 | C | T | missense_variant | MODERATE | Afu1g17200 | transcript | Coding | c.3401C>T | p.Ser1134Ph | 4316/15449 | 3401/14292 | 1277/4763 |
| | | Chr1_A_fumigatus_Af29 | 4692680 | C | T | synonymous_variant | LOW | Afu1g17200 | transcript | Coding | c.3831C>T | p.Ala1277Al | 4746/15449 | 3831/14292 | 1917/4763 |
| | | Chr1_A_fumigatus_Af29 | 4694600 | A | G | synonymous_variant | LOW | Afu1g17200 | transcript | Coding | c.5751A>G | p.Ala1917Al | 6666/15449 | 5751/14292 | 3171/4763 |
| | | Chr1_A_fumigatus_Af29 | 4698362 | C | A | synonymous_variant | LOW | Afu1g17200 | transcript | Coding | c.9513C>A | p.Pro3171Pr | 10428/1544 | 9513/14292 | 3200/4763 |
| | | Chr1_A_fumigatus_Af29 | 4698447 | G | A | missense_variant | MODERATE | Afu1g17200 | transcript | Coding | c.9598G>A | p.Gly3200Se | 10513/1544 | 9598/14292 | 3243/4763 |
| | | Chr1_A_fumigatus_Af29 | 4698576 | T | C | missense_variant | MODERATE | Afu1g17200 | transcript | Coding | c.9727T>C | p.Phe3243Le | 10642/1544 | 9727/14292 | 3273/4763 |
| | | Chr1_A_fumigatus_Af29 | 4698668 | G | A | synonymous_variant | LOW | Afu1g17200 | transcript | Coding | c.9819G>A | p.Leu3273Le | 10734/1544 | 9819/14292 | 3940/4763 |
| | | Chr1_A_fumigatus_Af29 | 4700669 | T | A | synonymous_variant | LOW | Afu1g17200 | transcript | Coding | c.11820T>A | p.Ala3940Al | 12735/1544 | 11820/1429 | 3979/4763 |
| | | Chr1_A_fumigatus_Af29 | 4700784 | T | G | missense_variant | MODERATE | Afu1g17200 | transcript | Coding | c.11935T>G | p.Ser3979Al | 12850/1544 | 11935/1429 | 4340/4763 |
| | | Chr1_A_fumigatus_Af29 | 4701868 | A | T | missense_variant | MODERATE | Afu1g17200 | transcript | Coding | c.13019A>T | p.Tyr4340Ph | 13934/1544 | 13019/1429 | 4741/4763 |
| | | Chr1_A_fumigatus_Af29 | 4703071 | G | A | missense_variant | MODERATE | Afu1g17200 | transcript | Coding | c.14222G>A | p.Gly4741Gl | 15137/1544 | 14222/14292 | |
| | | Chr1_A_fumigatus_Af29 | 4703730 | G | A | downstream_gene_variant | MODIFIER | Afu1g17200 | transcript | Coding | c.*589G>A | | | | |
| | | Chr1_A_fumigatus_Af29 | 4703980 | GCTGG | GCTGG | downstream_gene_variant | MODIFIER | Afu1g17200 | transcript | Coding | c.*852_*862delTGGAAAAAGTC | | | | |
| | P1MS | Chr1_A_fumigatus_Af29 | 4689600 | C | G | missense_variant | MODERATE | Afu1g17200 | transcript | Coding | c.751C>G | p.Pro251Ala | 1666/15449 | 751/14292 | 1131/4763 |
| | | Chr1_A_fumigatus_Af29 | 4692240 | A | G | missense_variant | MODERATE | Afu1g17200 | transcript | Coding | c.3391A>G | p.Asn1131A | 4306/15449 | 3391/14292 | 3200/4763 |
| *sidC* | | Chr1_A_fumigatus_Af29 | 4698447 | G | A | missense_variant | MODERATE | Afu1g17200 | transcript | Coding | c.9598G>A | p.Gly3200Se | 10513/1544 | 9598/14292 | 3243/4763 |
| | | Chr1_A_fumigatus_Af29 | 4698576 | T | C | missense_variant | MODERATE | Afu1g17200 | transcript | Coding | c.9727T>C | p.Phe3243Le | 10642/1544 | 9727/14292 | 3979/4763 |
| | | Chr1_A_fumigatus_Af29 | 4700784 | T | G | missense_variant | MODERATE | Afu1g17200 | transcript | Coding | c.11935T>G | p.Ser3979Al | 12850/1544 | 11935/1429 | 4340/4763 |
| | | Chr1_A_fumigatus_Af29 | 4701868 | A | T | missense_variant | MODERATE | Afu1g17200 | transcript | Coding | c.13019A>T | p.Tyr4340Ph | 13934/1544 | 13019/1429 | 4741/4763 |
| | | Chr1_A_fumigatus_Af29 | 4703071 | G | A | missense_variant | MODERATE | Afu1g17200 | transcript | Coding | c.14222G>A | p.Gly4741Gl | 15137/1544 | 14222/14292 | |
| | P2CS | Chr1_A_fumigatus_Af29 | 4689727 | C | T | missense_variant | MODERATE | Afu1g17200 | transcript | Coding | c.878C>T | p.Pro293Leu | 1793/15449 | 878/14292 | 1131/4763 |
| | | Chr1_A_fumigatus_Af29 | 4692240 | A | G | missense_variant | MODERATE | Afu1g17200 | transcript | Coding | c.3391A>G | p.Asn1131A | 4306/15449 | 3391/14292 | 1917/4763 |
| | | Chr1_A_fumigatus_Af29 | 4694600 | A | G | synonymous_variant | LOW | Afu1g17200 | transcript | Coding | c.5751A>G | p.Ala1917Al | 6666/15449 | 5751/14292 | 3200/4763 |
| | | Chr1_A_fumigatus_Af29 | 4698447 | G | A | missense_variant | MODERATE | Afu1g17200 | transcript | Coding | c.9598G>A | p.Gly3200Se | 10513/1544 | 9598/14292 | 3243/4763 |
| | | Chr1_A_fumigatus_Af29 | 4698576 | T | C | missense_variant | MODERATE | Afu1g17200 | transcript | Coding | c.9727T>C | p.Phe3243Le | 10642/1544 | 9727/14292 | 3781/4763 |
| | | Chr1_A_fumigatus_Af29 | 4700190 | C | T | missense_variant | MODERATE | Afu1g17200 | transcript | Coding | c.11341C>T | p.His3781Ty | 12256/1544 | 11341/1429 | 3979/4763 |
| | | Chr1_A_fumigatus_Af29 | 4700784 | T | G | missense_variant | MODERATE | Afu1g17200 | transcript | Coding | c.11935T>G | p.Ser3979Al | 12850/1544 | 11935/1429 | 4703/4763 |
| | | Chr1_A_fumigatus_Af29 | 4702957 | C | T | missense_variant | MODERATE | Afu1g17200 | transcript | Coding | c.14108C>T | p.Ser4703Le | 15023/1544 | 14108/1429 | 4741/4763 |
| | | Chr1_A_fumigatus_Af29 | 4703071 | G | A | missense_variant | MODERATE | Afu1g17200 | transcript | Coding | c.14222G>A | p.Gly4741Gl | 15137/1544 | 14222/14292 | |
| | | Chr1_A_fumigatus_Af29 | 4703360 | A | G | 3_prime_UTR_variant | MODIFIER | Afu1g17200 | transcript | Coding | c.*219A>G | | | | |
| | | Chr1_A_fumigatus_Af29 | 4703980 | GCTGG | GCTGG | downstream_gene_variant | MODIFIER | Afu1g17200 | transcript | Coding | c.*852_*862delTGGAAAAAGTC | | | | |
| | 08-19-02-30 D | Chr1_A_fumigatus_Af29 | 4690630 | T | C | missense_variant | MODERATE | Afu1g17200 | transcript | Coding | c.1781T>C | p.Leu594Ser | 2696/15449 | 1781/14292 | 1131/4763 |
| | | Chr1_A_fumigatus_Af29 | 4692240 | A | G | missense_variant | MODERATE | Afu1g17200 | transcript | Coding | c.3391A>G | p.Asn1131A | 4306/15449 | 3391/14292 | 1591/4763 |
| | | Chr1_A_fumigatus_Af29 | 4693620 | C | A | missense_variant | MODERATE | Afu1g17200 | transcript | Coding | c.4771C>A | p.Leu1591Ile | 5686/15449 | 4771/14292 | 1917/4763 |
| | | Chr1_A_fumigatus_Af29 | 4694600 | A | G | synonymous_variant | LOW | Afu1g17200 | transcript | Coding | c.5751A>G | p.Ala1917Al | 6666/15449 | 5751/14292 | 1933/4763 |
| | | Chr1_A_fumigatus_Af29 | 4694647 | T | C | missense_variant | MODERATE | Afu1g17200 | transcript | Coding | c.5798T>C | p.Met1933T | 6713/15449 | 5798/14292 | 3200/4763 |
| | | Chr1_A_fumigatus_Af29 | 4698447 | G | A | missense_variant | MODERATE | Afu1g17200 | transcript | Coding | c.9598G>A | p.Gly3200Se | 10513/1544 | 9598/14292 | 3243/4763 |
| | | Chr1_A_fumigatus_Af29 | 4698576 | T | C | missense_variant | MODERATE | Afu1g17200 | transcript | Coding | c.9727T>C | p.Phe3243Le | 10642/1544 | 9727/14292 | 3979/4763 |
| | | Chr1_A_fumigatus_Af29 | 4700784 | T | G | missense_variant | MODERATE | Afu1g17200 | transcript | Coding | c.11935T>G | p.Ser3979Al | 12850/1544 | 11935/1429 | 4433/4763 |
| | | Chr1_A_fumigatus_Af29 | 4702148 | C | T | synonymous_variant | LOW | Afu1g17200 | transcript | Coding | c.13299C>T | p.Asp4433A | 14214/1544 | 13299/1429 | 4741/4763 |
| | | Chr1_A_fumigatus_Af29 | 4703071 | G | A | missense_variant | MODERATE | Afu1g17200 | transcript | Coding | c.14222G>A | p.Gly4741Gl | 15137/1544 | 14222/14292 | |
| | | Chr1_A_fumigatus_Af29 | 4703980 | GCTGG | GCTGG | downstream_gene_variant | MODIFIER | Afu1g17200 | transcript | Coding | c.*852_*862delTGGAAAAAGTC | | | | |
| | 08-19-02-46 D | Chr1_A_fumigatus_Af29 | 4688032 | A | C | 5_prime_UTR_variant | MODIFIER | Afu1g17200 | transcript | Coding | c.-706A>C | | | | |
| | | Chr1_A_fumigatus_Af29 | 4688042 | T | C | 5_prime_UTR_variant | MODIFIER | Afu1g17200 | transcript | Coding | c.-696T>C | | | | 193/4763 |
| | | Chr1_A_fumigatus_Af29 | 4689426 | A | G | missense_variant | MODERATE | Afu1g17200 | transcript | Coding | c.577A>G | p.Ile193Val | 1492/15449 | 577/14292 | 398/4763 |
| | | Chr1_A_fumigatus_Af29 | 4690043 | A | T | synonymous_variant | LOW | Afu1g17200 | transcript | Coding | c.1194A>T | p.Leu398Leu | 2109/15449 | 1194/14292 | 470/4763 |
| | | Chr1_A_fumigatus_Af29 | 4690259 | A | G | synonymous_variant | LOW | Afu1g17200 | transcript | Coding | c.1410A>G | p.Thr470Thr | 2325/15449 | 1410/14292 | 523/4763 |
| | | Chr1_A_fumigatus_Af29 | 4690418 | C | G | missense_variant | MODERATE | Afu1g17200 | transcript | Coding | c.1569C>G | p.Ser523Arg | 2484/15449 | 1569/14292 | 771/4763 |
| | | Chr1_A_fumigatus_Af29 | 4691160 | G | A | missense_variant | MODERATE | Afu1g17200 | transcript | Coding | c.2311G>A | p.Ala771Thr | 3226/15449 | 2311/14292 | 1131/4763 |
| | | Chr1_A_fumigatus_Af29 | 4692240 | A | G | missense_variant | MODERATE | Afu1g17200 | transcript | Coding | c.3391A>G | p.Asn1131A | 4306/15449 | 3391/14292 | 1274/4763 |
| | | Chr1_A_fumigatus_Af29 | 4692669 | A | G | missense_variant | MODERATE | Afu1g17200 | transcript | Coding | c.3820A>G | p.Ile1274Val | 4735/15449 | 3820/14292 | 1917/4763 |
| | | Chr1_A_fumigatus_Af29 | 4694600 | A | G | synonymous_variant | LOW | Afu1g17200 | transcript | Coding | c.5751A>G | p.Ala1917Al | 6666/15449 | 5751/14292 | 2161/4763 |
| | | Chr1_A_fumigatus_Af29 | 4695332 | C | T | synonymous_variant | LOW | Afu1g17200 | transcript | Coding | c.6483C>T | p.Phe2161Pl | 7398/15449 | 6483/14292 | 2242/4763 |
| | | Chr1_A_fumigatus_Af29 | 4695575 | G | A | synonymous_variant | LOW | Afu1g17200 | transcript | Coding | c.6726G>A | p.Arg2242Ar | 7641/15449 | 6726/14292 | 2392/4763 |
| | | Chr1_A_fumigatus_Af29 | 4696023 | A | G | missense_variant | MODERATE | Afu1g17200 | transcript | Coding | c.7174A>G | p.Lys2392Gl | 8089/15449 | 7174/14292 | 3200/4763 |
| | | Chr1_A_fumigatus_Af29 | 4698447 | G | A | missense_variant | MODERATE | Afu1g17200 | transcript | Coding | c.9598G>A | p.Gly3200Se | 10513/1544 | 9598/14292 | 3234/4763 |

| | | | | | | | | | | | | | |
|---|---|---|---|---|---|---|---|---|---|---|---|---|---|
| Chr1_A_fumigatus_Af29: | 4698551 | C | G | synonymous_variant | LOW | Afu1g17200 | transcript | Coding | c.9702C>G | p.Leu3234Le | 10617/1544 | 9702/14292 | 3243/4763 |
| Chr1_A_fumigatus_Af29: | 4698576 | T | C | missense_variant | MODERATE | Afu1g17200 | transcript | Coding | c.9727T>C | p.Phe3243Le | 10642/1544 | 9727/14292 | 3257/4763 |
| Chr1_A_fumigatus_Af29: | 4698618 | C | A | missense_variant | MODERATE | Afu1g17200 | transcript | Coding | c.9769C>A | p.Gln3257Ly | 10684/1544 | 9769/14292 | 3940/4763 |
| Chr1_A_fumigatus_Af29: | 4700669 | T | A | synonymous_variant | LOW | Afu1g17200 | transcript | Coding | c.11820T>A | p.Ala3940Al | 12735/1544 | 11820/1429 | 3979/4763 |
| Chr1_A_fumigatus_Af29: | 4700784 | T | G | missense_variant | MODERATE | Afu1g17200 | transcript | Coding | c.11935T>G | p.Ser3979Al | 12850/1544 | 11935/1429 | 4356/4763 |
| Chr1_A_fumigatus_Af29: | 4701916 | A | C | missense_variant | MODERATE | Afu1g17200 | transcript | Coding | c.13067A>C | p.Asn4356Th | 13982/1544 | 13067/1429 | 4475/4763 |
| Chr1_A_fumigatus_Af29: | 4702274 | T | C | synonymous_variant | LOW | Afu1g17200 | transcript | Coding | c.13425T>C | p.Ser4475Se | 14340/1544 | 13425/1429 | 4741/4763 |
| Chr1_A_fumigatus_Af29: | 4703071 | G | A | missense_variant | MODERATE | Afu1g17200 | transcript | Coding | c.14222G>A | p.Gly4741Gl | 15137/1544 | 14222/14292 | |
| Chr1_A_fumigatus_Af29: | 4703730 | G | A | downstream_gene_variant | MODIFIER | Afu1g17200 | transcript | Coding | c.*589G>A | | | | |
| Chr1_A_fumigatus_Af29: | 4703980 | GCTGG | GCTGC | downstream_gene_variant | MODIFIER | Afu1g17200 | transcript | Coding | c.*852_*862delTGGAAAAGTC | | | | |
